# Generation of tumor organoids and cancer-associated fibroblasts from minimal pancreatic cancer biopsies enables investigation of tumor-stroma interactions

**DOI:** 10.64898/2026.11.25.769316

**Authors:** Emilie Jaune-Pons, Fatemeh Ardakani Behjati, Maria Nica, Courtney Brooks, Jennifer Campbell, Jennifer L. Gorman, Noor Shakfa, Naila Abid, Phoebe Ji, Le Tran Phuc Khoa, Peter Sajjakunukit, Gavin Goebel, Domenic De Stasi, Leah J.J. McDonald, Dami Otukoya, Costas A. Lyssiotis, Matthew Cecchini, Kenneth A. Leslie, Stephen A. Welch, Hartland W. Jackson, Nadeem Hussain, Michael Sey, Brian M. Yan, Christopher L. Pin

## Abstract

Pancreatic ductal adenocarcinoma (PDAC) remains among the deadliest forms of cancer, accounting for the third leading cause of cancer-related deaths in North America. A critical factor in this poor outcome is the inability to effectively treat patients not eligible for surgery. The development of patient-derived organoids (PDOs) from tumors has shown promise as a personalized approach for treating PDAC. In the case of PDAC, PDOs need to be generated immediately upon diagnosis due to the rapid mortality observed in these patients. In addition, the tumor microenvironment, a significant factor in PDAC, needs to be considered. To address these two issues, we developed an approach to isolate both cancer and non-cancer-cells of PDAC tumors through endoscopic ultrasound at the time of initial diagnosis. PDOs and cancer-associated fibroblast (CAF) cultures retain patient-specific heterogeneity related to the *in vivo* tumor phenotype with PDOs showing two distinct and stable phenotypic and molecular profiles. These subtypes were affected by introducing conditioned media from primary CAF cultures, which promoted transcriptomic and epigenetic changes in PDOs reflective of a more resistant phenotype. This study identifies approaches for rapid establishment of multiple tumor compartments *ex vivo* and highlights the importance in considering the tumor microenvironment when assessing the effects of chemotherapeutic responses in patients.

## Introduction

Pancreatic ductal adenocarcinoma (PDAC) has the worst prognosis of any of the major cancers, with a 5-year survival rate of ∼13% worldwide.^1^ This is the result of late diagnosis, heterogeneity between and within patients, and ineffective responses to chemotherapy, the main therapeutic option for 80% of PDAC patients. While there is significant promise in the recent development of KRAS inhibitors such as daraxonrasib, the two main therapies for PDAC are FOLFIRINOX (a combination of 5-fluorouracil, irinotecan, leucovorin and oxaliplatin) or Gemcitabine combined with nab-paclitaxel (i.e. Abraxane^®^).^2^ Part of the inefficiency underlying chemotherapy treatment is the lack of quantifiable markers that indicate the responsiveness of each patient. In addition, while novel therapies can be suggested as second line therapies, these options are limited by a lack of understanding on the biology of each patient’s tumor. This includes non-cancer cells of the tumor, which impact the tumor microenvironment and affect cancer cell phenotypes and resistance.^3,4^

In the past few years, the isolation and characterization of patient-derived organoids (PDO) from PDAC tumors have been suggested as an approach for personalized medicine. PDOs provide unique molecular information specific to the individual and retain important molecular, epigenetic, and drug response effects that can be used to inform treatment.^5–7^ However, obtaining and characterizing PDO’s at the earliest time point in PDAC is critical for informing treatment, given the rapid progression observed for this cancer. Additionally, the tumor microenvironment (TME) needs to be incorporated into this approach as studies suggest accurate PDO responses to therapy may require both cancer and non-cancer cells to mimic patient responses.^8,9^ The TME contains numerous cell types including cancer-associated fibroblasts (CAFs), immune cells, and neuronal cells, as well as a significant stromal component, which contributes to a hypovascular and hypoxic environment in PDAC.^10^ In particular, CAFs play an important role in shaping the TME, affecting cancer cell phenotypes and promoting resistance.^11–14^ Like tumor cells, CAFs are heterogeneous between and within patient tumors. Several different CAF subtypes have been described in the literature with most studies defining myofibroblast (my) CAFs, inflammatory (i) CAFs, and antigen presenting (ap) CAFs.^11,15^ Since CAFs affect tumor biology, their contributions to patient-specific resistance need to be considered.

To address these issues, we developed a tissue bank that captures cancer and non-cancer cell components from the earliest time point in a patient’s therapeutic odyssey. We obtained fine needle biopsies from patients undergoing diagnostic endoscopic ultrasound (EUS) for newly identified pancreatic masses. We processed these biopsies to generate PDO’s of cancer cells in 3D, and isolated non-cancer cells enriched for CAFs in 2D. Using this novel biobank of samples, we showed EUS-derived PDOs and CAFs retain their *in vivo* phenotype and show significant heterogeneity between patients. PDOs exhibit a range of genetic mutations and gene expression profiles and CAFs retain heterogeneity for several passages. CAF samples enriched for iCAFs promote transcriptional and epigenetic changes in PDOs leading to a more resistant phenotype. These findings highlight the importance of retaining TME components when assessing PDO potential for identifying therapeutic sensitivity.

## Methodology

All reagents were purchased from Wisent, Montreal, QC except where noted.

### EUS and Fine Needle Biopsy

Patients with a suspected or known diagnosis of PDAC, based on clinical and radiological criteria, were consented into DERIVE (Determination of response to therapy in individual patients, REB# 113362 approved by the London Health Sciences Centre and Western Ethics committee) from 2021-2025. Upon consent, after an adequate sample was obtained for clinical purposes, additional 1-2 fine needle biopsies (FNB) was obtained using either 22-gauge SharkCore (Medtronic, Brampton, ON) or Cook EchoTip ClearCore needles (Cook Medical, Bloomington, IN). In the event of metastatic disease, the specimen for DERIVE was obtained from the primary tumor within the pancreas. Biopsies were immediately placed in human wash media and transferred to core research laboratories where, if sufficient material was obtained, a small portion of the core was removed and processed for formalin-fixed, paraffin-embedded (FFPE). The remaining tissue was used to prepare PDOs and CAF-enriched cultures.

### PDOs and CAFs isolation

Over the course of this study, we used modified the protocol for organoid and CAF isolation numerous times to increase efficiency. Many of the previous methodologies published for isolating organoids were inaccurate or left out critical details. For example, we attempted to establish 3D cultures by initially culturing on BME layers, but over time switched to growing cells in 50 µL domes of BME. An extensive protocol (see Supplemental Methodology) is provided that resulted in the most successful isolation and establishment of PDOs and CAFs from a single FNB.

For passaging or experiments, media was removed and PDOs incubated for 1 hour at 4°C in 800 µL of ice-cold cell recovery solution (Corning, #354253), then resuspended, centrifuged and pellets incubated into Accutase supplemented with 10.6 µM Y-27632 and 10 µg/mL DNase I (Sigma, #10104159001) for 45 min at 37°C under agitation. Digested PDOs were centrifuged and rinsed with human wash media + 0.1% BSA and grown in domes.

### RNA and DNA isolation and analysis from PDOs

PDOs were plated in domes of BME. After 14 days, media was removed and 300 µL of cold cell recovery solution added and incubated for 1 hour at 4°C. Organoids from three domes were combined and centrifuged for 5 minutes at 2,000 rpm and rinsed twice with cold PBS. RNA was extracted from organoid pellets following the RNeasy plus mini kit protocol (Qiagen, #74134). RNAs were eluted in 30 µL of Rnase-free water and RNA concentration measured by Qubit (Invitrogen, #Q32852). DNA from organoids pellet was extracted following the Dneasy Blood and tissue kit (Qiagen, #69504). DNAs were eluted in 30 µL of Buffer AE and DNA concentration was measured by Qubit (Invitrogen, #Q32851). Chromatin was extracted from organoids using the ATAC-Seq Kit (Active Motif, #53150). Chromatin was eluted in 20 µL of DNA purification elution buffer and concentration was measured by Qubit (Invitrogen, #Q32851).

### RNA-Sequencing

Raw RNA-seq reads in FASQ format were processed using Trimmomatic v0.39 in paired-end mode with the Phred33 quality encoding option. Adapter trimming was performed with the NexteraPE-PE adapter set using the following parameters: *ILLUMINACLIP:NexteraPE-PE.fa:2:30:10 LEADING:3 TRAILING:3 SLIDINGWINDOW:4:15 MINLEN:36*. Trimmed reads were aligned to the human reference genome (GRCh38.p14) using STAR v2.7.11a with default parameters. The resulting alignment files were processed with samtools v1.19.2 to generate sorted and indexed BAM files. Gene-level read counts were then quantified using featureCounts v2.0.8, based on the Ensembl *Homo_sapiens.GRCh38.112.gtf* annotation. Genes were filtered out by discarding those with counts per million (CPM) values below the threshold corresponding to 10 raw read counts. Genes were retained if their CPM exceeded this threshold in at least two samples, and transcripts per million (TPM) computed for the remaining genes to normalized expression levels across samples.

For principal component analysis (PCA) analysis of PDOs, we performed PCA on filtered and normalized log,[TPM + 1] values from organoids treated with human feeding media (HFM) using the *prcomp* function in R with default settings. Top 50 most variable genes were identified based on the variance of log,[TPM + 1] values across samples treated with HFM. These genes were plotted as a heatmap. Additionally, marker genes for basal-like and classical tumor subtypes were curated from the literature. Classical and basal markers have been determined according to the following studies^16–19^. Log, TPM values were visualized as a heatmap to examine expression patterns across samples grouped by tumor subtype.

For the differential expression from PDOs-treated with CAF conditioned media, DESeq2 v1.44.0 analysis was performed on classical samples to identify genes differentially expressed in response to environmental changes in organoids. We contrasted iCAF vs. HFM and myCAF vs. HFM conditions, selecting differentially expressed genes (DEGs) with adjusted P ≤ 0.05 and |log, fold change| ≥ 1. The resulting DEG sets were subsequently used to perform Reactome pathway enrichment analysis with a false discovery rate (FDR) threshold ≤ 0.05, identifying significantly enriched pathways. For differential expression between CAF subtypes, DEGs between iCAF and myCAF subtypes were identified using DESeq2 v1.44.0 on raw read counts from CAF RNA-seq samples. The design matrix included subtypes (iCAF vs. myCAF) as the primary factor.

### Flow cytometry

All analysis and experiments using patient-derived CAFs were carried out within nine passages of initial isolation. To characterize CAF phenotypes, 2D cultures were grown to 80% confluence, then detached with diluted Accutase (50% PBS + 5 mM EDTA) and resuspended in 1 mL cold FACS (fluorescence-activated cell sorting) buffer (PBS, 5% BSA, 5 mM EDTA). Cells were incubated at 4°C in the dark with conjugated antibodies described in Supplementary Table 1. Following incubation, cells were centrifuged for 5 minutes at 2,000 rpm at 4°C, rinsed and fixed in 2% PFA at 4°C for 10 min. Following a second centrifugation, cells were resuspended in 400 µL of FACS buffer and analyzed on a BD Biosciences LSR-II flow cytometer.

### Histological analysis

A portion of each EUS biopsy was prepared for histology, or pathological assessment was performed on additional biopsies from EUS. To embed PDOs, isolated organoids were centrifuged at 410 rcf for 3 min at 4°C then fixed in 10% formalin and incubated for 15 min at room temperature. Samples were washed 3x in PBS and then incubated in 100% ethanol for 20 min. Ethanol was removed and PDOs resuspended in HistoGel (Epredia, #HG-4000-012). Tissues were sectioned to 5 µm thickness for histology and IHC or IF. For general histology, Hematoxylin and Eosin staining was performed. Sections were dehydrated, mounted with Permount (Fisher Chemical, #SP15-100) and analyzed using a Leica microscope (Leica DM5500 B).

For immunofluorescence, antigen retrieval was performed by incubating sections in citrate solution for 45 minutes at 90°C, then a further 10 minutes at room temperature in 50 mM NH_4_Cl. To permeabilize tissue, sections were incubated with 0.2% Triton-X in PBS. Samples were rinsed and covered with blocking solution (5% serum of secondary antibody species + 0.2% Triton in PBS) for 1 hour. Sections were then incubated overnight at 4°C with primary antibody (Supplementary Table 1). The following day, slides were rinsed and incubated in secondary antibody coupled with TRITC or FITC (Jackson ImmunoResearch, 1/200) for 1 hour at room temperature. Sections were rinsed and mounted with Permount and analyzed with a Leica microscope (Nikon Eclipse Ti2).

### Confirmation of KRAS mutations

To confirm *KRAS* mutations, PCR was performed on genomic DNA using primers flanking exon 2 of the KRAS gene (NM_004985). Primers used for amplification were the following primers:

*KRAS* forward (KRAS12f) 5’ GAAGTACAGTTCATTACGATACACG 3’

*KRAS* reverse (KRAS12r) 5’ CATTTCAGATAACTTAACTTTCAGCAT 3’

PCR was performed using the PCR kit from Takara (Clontech, #638500) following the CloneAmp kit instructions for 35 cycles and an annealing temperature of 58°C. Amplified samples were purified using the QIAquick PCR purification kit (Qiagen, # 28104) and elude with 40 µL of EB buffer. Sanger sequencing was performed using KRAS12f and KRAS12r respectively.

### Imaging Mass Cytometry analysis

Formalin-fixed, paraffin-embedded (FFPE) sections were processed as previously described^55^ and incubated overnight at 4°C with titrated metal-labelled primary antibodies (Supplementary Table 2), with secondary amplification where required. Slides were washed with TBS, counterstained with an iridium intercalator (500 nM) for 5 min, and rinsed 3x in TBS, once with deionized water, dried, and acquired Imaging Mass Cytometry (IMC).

Antibody performance was optimized in control tissues including human tonsil, spleen, appendix, and PDAC. Slide preparation for optimization followed the same protocol as described above. Antibody specificity and marker co-localization were confirmed by IF and nuclei counterstained with DAPI. Validated antibodies were conjugated to metal isotopes using the Maxpar X8 Multi-Metal Antibody Labelling Kit (Standard BioTools) and stored in Candor Antibody Stabilizer (Candor Biosciences) at 4°C until use.

IMC data were acquired using the Hyperion XTi Imaging System (Standard BioTools) at 1 µm resolution, in a raster scan pattern of 800 Hz. Acquisition order was randomized and blinded to patient identifiers. Raw data were collected using the Standard BioTools acquisition software. Images were inspected using Rakaia (https://rakaia.io/)^20^ and segmented into single cells using Mesmer (DeepCell^21^) based on DNA, CD45, pan-cytokeratin and vimentin signals. For each region of interest (ROI), single-cell masks and expression matrices were generated, with marker intensities capped at the 99.9^th^ percentile and normalized to the maximum value for each marker across all samples.

ROIs (n = 273 total) were quality controlled using (1) number of segmented objects (cellularity) per mcd file, with low cellularity ROIs excluded while retaining enough ROIs per patient to capture at least 75% of the total cells, (2) marker-expression threshold filtering based on predefined expression thresholds for key markers, and (3) manual image review. Specifically, ROIs were flagged for visual inspection based on high SMA expression and/or low DNA and excluded if the staining quality was insufficient. Final dataset (273 ROIs) quality was further assessed in Rakaia and by review of PhenoGraph clustering results.

Dimensionality reduction was performed by PCA, followed by PhenoGraph clustering in R CAFs were annotated using SMA, PDGFRα and canonical markers for apCAFs (HLA-DR^+^ or CD74^+^), myCAFs (SMA^+^ with higher PDGFRβ), and iCAFs (PDGFRα^+^ or CXCL12^+^). Where CXCL12 was elevated in PDGFRβ+ clusters, SMA was used to resolve assignments, with SMA^+^ cells classified as myCAFs and SMA– cells as iCAFs. All visualization and statistical analyses were performed in R using the ggplot2 package and appropriate statistical tests.

### Analysis of ATAC-Sequencing

Raw ATAC-seq reads in FASQ format were processed using Trimmomatic v0.39 in paired-end mode with the Phred33 quality encoding option. Adapter trimming was performed with the NexteraPE-PE adapter set using the following parameters:

### ILLUMINACLIP:NexteraPE-PE.fa:2:30:10 LEADING:3 TRAILING:3 SLIDINGWINDOW:4:15 MINLEN:3C

Trimmed reads were aligned to the human reference genome (GRCh38.p14) using STAR v2.7.11a with default parameters. The resulting alignment files were processed with samtools v1.19.2 to generate sorted and indexed BAM files. Genome-wide coverage tracks were generated using bedtools v2.31.1. For each sample, per-base read coverage was computed from the sorted BAM files in BedGraph format using the *genomecov* function. These procedures yielded genome-wide coverage profiles, which were used for data visualization and downstream comparative analyses. Transcription start site (TSS) coordinates for all annotated genes were obtained from the Ensembl *Homo_sapiens.GRCh38.112.gtf* annotation.

For each gene, a 2 kb genomic window centered on the TSS (±1 kb) was defined using bedtools v2.31.1. This was achieved by extending each TSS region symmetrically upstream and downstream relative to the reference genome boundaries. To enable higher-resolution analysis, each 2 kb window was subsequently partitioned into 100 bp non-overlapping bins using the *makewindows* function in bedtools. ATAC-seq signal intensity within each bin was quantified by computing the total number of reads overlapping each interval with the *map* function of bedtools, using parameters that summed the read counts while assigning a value of zero to bins with no coverage. This procedure generated a matrix of gene-by-bin values (20 bins per gene), representing chromatin accessibility profiles across TSS-proximal regions with profiles of minus-strand genes mirrored to maintain transcriptional orientation consistency. The resulting matrix was used to generate heatmaps of chromatin accessibility (Figure 6B) for selected patient-derived organoids (PDOs). A subset of the matrix was further used to display chromatin profiles over a 1 kb window centered on the TSS (±0.5 kb) for selected genes in organoids treated with iCAF, myCAF, or HFM.

### Metabolomic analysis

LC-MS analyses were performed using an Agilent Technologies Triple Quad 6470 LC-MS/MS system as previously described^22^. The setup previously described was utilized for negative ion-mode metabolites. An additional method was used with the same hardware configuration to detect positive ion-mode metabolites. In brief, a Waters Acquity UPLC BEH TSS C18 column (2.1 × 100mm, 1.7µm) with ACQUILITY UPLC C181.7µm VanGurad Pre-Colum was used with mobile phase (A) consisting of 0.1% formic acid in water; mobile phase (B) consisting of 0.1% formic acid in acetonitrile. Using a gradient program, mobile phase (B) was held at 1% for 3 minutes, increased to 6% in 12 minutes, then to 15% B in 15 minutes, then to 99% B in 17 minutes and held for 2 min before going to the initial condition at 19.1 min and held for 5 minutes. The column was at 40 °C and 3 µl of sample was injected into the LC-MS with a flow rate of 0.2 ml/minute. Key parameters of electrospray ionization (ESI) were gas temperature = 275°C, gas flow = 14 l/min, nebulizer at 20 psi, SheathGasHeater = 250°C, SheathGasFlow = 11 l/min, Capillary = 3000 V. MS: Delta EMV 200 V, Cycle Time 500 ms, Cell Acc 4 V.

Metabolite dMRMs were optimized through MassHunter Workstation Optimizer and added to the acquisition method. Data for both ion-mode detection methods was pre-processed with Agilent MassHunter Quantitative Analysis v10 to ensure accurate area-under-the-curve (AUC) chromatogram integration. The raw data was further processed in Microsoft Excel by median-centering and conducting Student T-Tests for significance.

## Results

### Patient-derived cancer lines were developed using 2D and 3D approaches from EUS sampling

Patients were enrolled into DERIVE (Determination of Response to Therapy in Individual Patient), a local-clinical database capturing individuals diagnosed with gastrointestinal pathologies. From DERIVE, patients with suspected or known PDAC pathology undergoing confirmatory diagnostic endoscopic ultrasound (EUS) were consented for an additional biopsy. Individuals undergoing neoadjuvant therapy were excluded from the study. A total of 202 individuals were recruited over five years. Of these biopsies, 15 were discarded due to low quality (i.e. excessive blood), 21 due to processing error and 25 due to contamination. The final cohort consisted of 122 individuals with PDAC, 4 with pancreatic neuroendocrine tumours, 5 with pancreatitis and 10 with other cancer subtypes (**Table 1**).

**Table 1:** Summary of EUS-derived cultures.

| **Characteristic** | **Total (%)** | **Male** | **Female** |
|---|---|---|---|
| #Enrolled | 202 | 113 | 89 |
| Biopsies excluded due to: |  |  |  |
| *Processing error* | 21 | 14 | 7 |
| *low quality FNB* | 15 | 10 | 5 |
| *contamination* | 25 | 12 | 13 |
| *Total* | 61 | 36 | 25 |
| Usable biopsies 3D | 141 | 77 | 64 |
| Usable biopsies 2D* | 103 | 50 | 53 |
| Pathology of usable biopsies |  |  |  |
| *PDAC* | 122 (86.5)** | 51 | 60 |
| *PNET* | 4 (2.8)** | 4 | 0 |
| *Other tumour* | 10 (7.1)** | 6 | 4 |
| *Pancreatitis* | 5 (3.6)** | 5 | 0 |
| Organoids visible (for all biopsies) | 124 (87.9)** | 76 | 59 |
| Organoids >P3 | 64 (52.5)# | 44 | 31 |
| Organoids >P5 | 31 (25.4)# | 11 | 20 |
| Line established*** | 15 (12.1)# | 3 | 12 |

<sup>\*</sup>2D cultures were not attempted for every EUS sample; \*\*percentage of the number of usable biopsies; \*\*\*all lines established are confirmed pancreatic ductal adenocarcinoma; #percentage of success relative to the number of PDAC samples (i.e. 122)

87.9% of the cultures (125/141) developed multi-cellular structures and were expanded to at least one passage (**Supplemental Figure 1A**). This success decreased to 52.5% percent by passage (P) 3 (64/141) and to 25.4% (31/141) by P5. While reduction in PDOs reaching P5 may be from the initial isolation of normal tissue, DNA sequence analysis confirmed several unsuccessful PDOs contained cancer cells based on the presence of *KRAS* mutations. From the original 122 samples, 15 resulted in established PDO lines (12.3% success; **Table 2**). PDOs could be classified into three groups based on morphology, similar to previous studies.^23^ LHSC04, 137 and 143 showed a gland-like (GL) phenotype with a simple epithelial layer (**Figure 1A**) surrounding a central lumen. LHSC107 and 134 showed a densely proliferative (DP) phenotype with stratified epithelium and multiple lumens, while LHSC124 exhibited both GL and DP phenotypes (**Figure 1A**).

**Figure 1.**
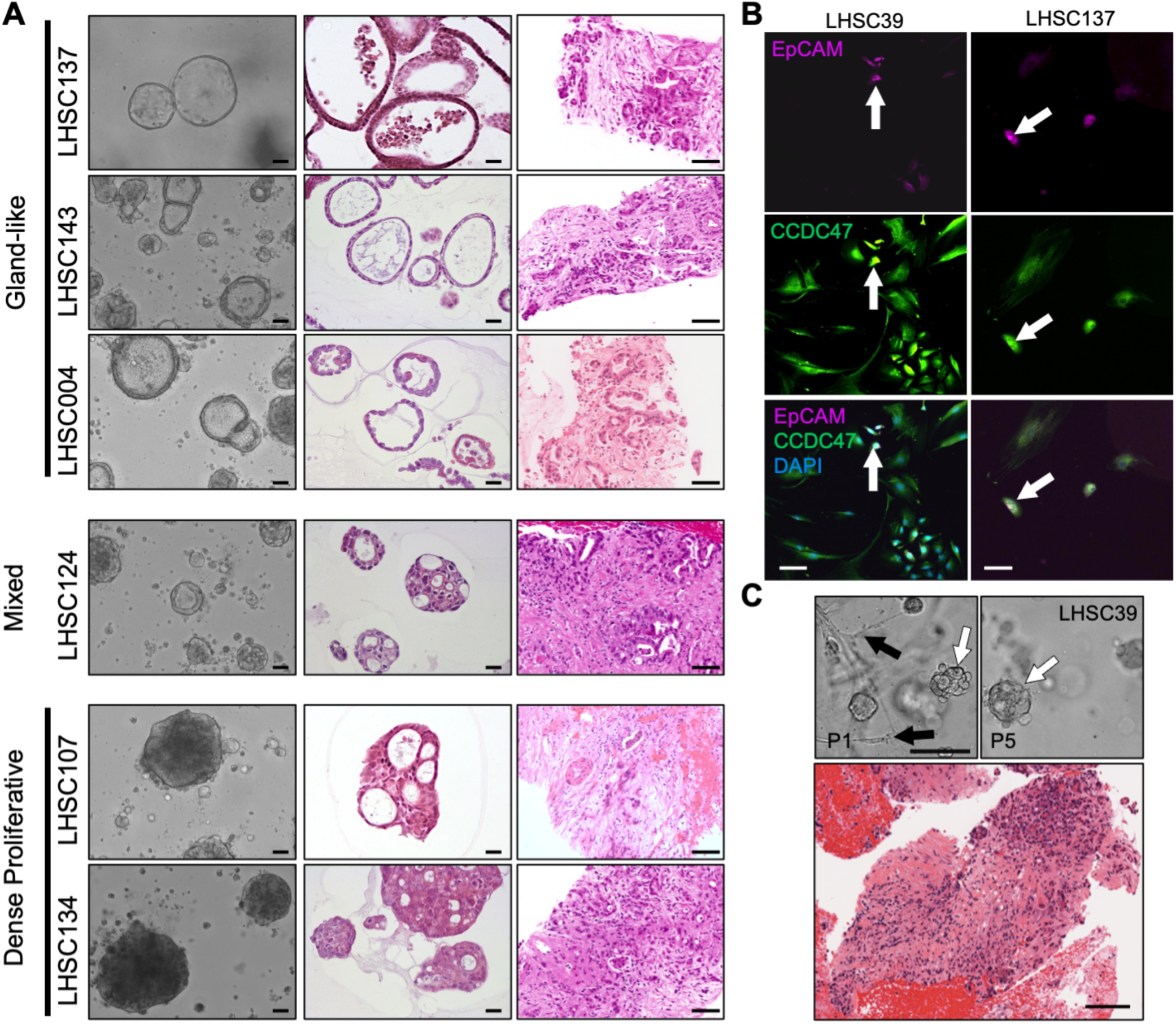
**(A)** Patient derived organoids (PDO) brightfield (Scale bar = 50µm) and H&E (Scale bar = 50µm) showing morphological differences between gland-like (GL), densely proliferative (DP) or mixed morphology. Right images are showing corresponding patient H&E (scale bar = 50 µm), presenting similar morphology. **(B)** Immunofluorescence on 2D cells isolated from patient samples identifying cancer cells with EpCAM marker (purple), all the cells with αCCDC47 marker (green) and nuclei with Dapi (blue; scale bar = 30 µm. White arrows are pointing out to cancer cells. **(C)** PDO from LHSC39 isolated in 2D brightfield (scale bar = 100 µm) showing a DP morphology at passage 1 (P1) and 5 (P5). White arrows are pointing out cancer cells. We can observe CAFs only at P1, highlighted by the black arrow. Bottom images are showing corresponding patient H&E, presenting similar morphology (scale bar = 50 µm).

**Table 2.** Characteristics of Organoid Lines.

| **EUS** | **Sex** | **PurIST<sup>1</sup>** | **KRAS** | **P53** | **SMAD4** | **Growth rate (days)** | **Sensitivity to GEM** |
|---|---|---|---|---|---|---|---|
| LHSC04 | F | 0.007 | G12D | S250Dup | R361H | 9.41 | 23.19 nM |
| LHSC22 | M | ND | G12D | G279E | WT | ND | ND |
| LHSC27 | NA | ND | G12D | A161T | ND | 10.02 | 4.38 nM |
| LHSC39* | M | 0.991 | G12R | WT | WT | ND | ND |
| LHSC107 | F | 0.427 | G12D | 175 WT | ND | 8.34 | 24.5 nM |
| LHSC124 | M | 0.049 | G12V | 175 WT | ND | 12.69 | 27.26 nM |
| LHSC134 | F | 0.228 | G13D | N175H | ND | 10.98 | 14.15 nM |
| LHSC137 | F | 0.003 | G12V | P278S | Arg361C | 7.78 | 19.85 nM |
| LHSC143 | F | 0.008 | G12D |  | D537T* | 9.98 | 23.26 nM |
| LHSC190 | F |  |  |  |  |  |  |
| LHSC194 | F |  | G12D |  |  |  |  |
| LHSC203** | F |  | G12D |  |  |  |  |
| LHSC204 | M |  | - |  |  |  |  |
| LHSC208 | F |  | G61L |  |  |  |  |
| LHSC213 | F |  | - |  |  |  |  |
| LHSC214 | F |  | G12V |  |  |  |  |

ND – Not determined; 1 - PurIST scores is a ratio score where classical PDOs <0.50;
\*line developed from CAF sample and grown in 2D; \*\*organoids developed following growth in 2D for 3 passages than transferred to 3D

To determine if the PDO lines conserve the same mutations as patients, we identified *KRAS* mutations through PCR/Sanger sequencing or Oncomine sequencing. Our patient cohort showed a range of *KRAS* mutations. The most common *KRAS* mutations in PDAC, including *KRAS^G12D^*, *KRAS^G12V^*, and *KRAS^G12R^* were identified in PDOs, as well as more rare mutations, such as *KRAS^G13D^*, indicating PDOs could be established from most mutant *KRAS* backgrounds.

As an alternative approach for obtaining patient-derived cancer lines, the remaining cell slurry was plated directly on tissue culture plates. These cultures developed as monolayers with cells exhibiting heterogeneity in morphology (**Supplemental Figure 1C-E**). Co-IF for the ubiquitously-expressed CCDC47 and EpCAM confirmed the presence of putative cancer cells in ∼9% of monolayer cultures examined (**Figure 1B**). Amplification and sequencing for *KRAS* confirmed oncogenic mutations in samples containing EpCAM^+^ cells (**Table 2, Appendix 1**). To determine if these cultures could be propagated into 3D organoids, LHSC39 cultures were seeded in matrigel. The resulting organoids took on a DP phenotype (**Figure 1C**) and could be grown over several passages. The resulting organoids showed reduced viability and growth upon passaging, with fibroblasts surviving only a single passage, suggesting cells within the co-cultures supported continued cancer cell growth.

### PDOs represent a range of morphological subtypes with a classical molecular profile

To characterize PDOs in more detail, histological (**Figure 1**) and molecular analysis (**Figure 2**) were performed on six PDO lines. To determine if *ex vivo* phenotypes were consistent with the *in vivo* makeup of a patient’s tumor, the histology of core biopsies from which these samples were obtained was examined. While some heterogeneity was observed in pathological samples, the predominant PDAC morphology (i.e. well-differentiated vs. poorly differentiated) was consistent with GL vs DP organoid histology, respectively (**Figure 1A**). Histological analysis of LHSC39 showed a poorly differentiated phenotype consistent with its DP phenotype in culture (**Figure 1C**)

**Figure 2.**
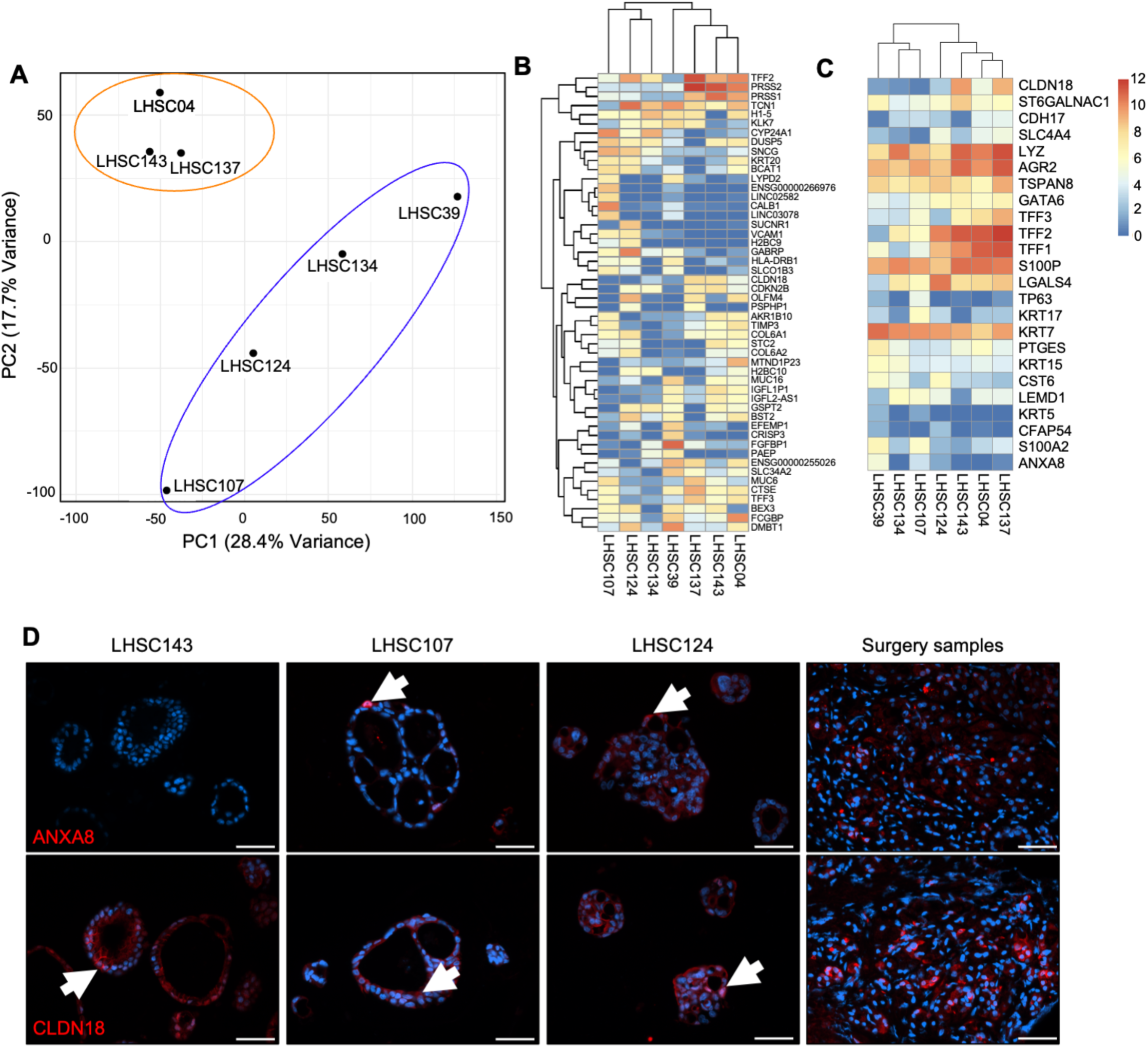
**(A)** Principal component analysis of seven PDOs showing GL-PDOs grouping together (orange circle) and mixed or DP-PDOs well separated from GL-PDOs (blue circle). **(B)** Heatmap of top 50 most variable genes across seven PDOs. This unsupervised hierarchical clustering shows mixed and DP-PDOs clustering away from GL-PDOs. **(C)** Heatmap of classical and basal genes in seven PDOs. Classical gene expressions like CLDN18, LYZ or TFF1/2 are increased in GL-PDOs versus mixed or DP-PDOs. **(D)** Immunofluorescence staining of ANXA8 (first line) or CLDN18 (second line) in red and of nuclei in blue by DAPI in three different organoids. ANXA8 is a basal marker and show higher staining in mixed or DP-PDOs. CLDN18 is a classical marker and show increased staining in GL and mixed-PDOs. Surgery samples are used as control (scale bar =50 µM).

Next, bulk RNA-seq was performed on PDOs. PCA grouped PDOs consistent with morphological appearances (**Figure 2A**), especially GL PDOs. PuRIST analysis, which defines classical or basal-like subtypes of PDAC based on gene expression signatures (**Table 2**)^24^ was performed. All PDOs showed PuRIST scores <0.5 except LHSC39, which had a PuRIST score of 0.99, indicating a basal-like phenotype. Interestingly, the DP PDOs, LHSC107 and LHSC134, showed PuRIST ratios much closer to a basal-like profile (>0.2), consistent with basal-like histological features (**Table 2**). The GL PDOs had PuRIST scores <0.01 while the mixed LHSC124 PDO had an intermediate PuRIST score (0.049).

Next, top 50 DEGs that explained segregation of the PDOs was identified. Hierarchical clustering aligned GL PDOs away from DP PDOs. Similar to phenotypic analysis, LHSC124 was intermediate between the two groups while LHSC39 was more divergent from GL PDOs (**Figure 2B**). Analysis of genes linked to basal-like or classical PDAC phenotypes confirmed higher levels of classical markers (*GATA6*, *CLDC18*, *LYZ* and *TFF1/2*) in the GL organoids. Genes linked to the basal-like state (*ANXA8, S100A2* and *KRT17*) showed variable expression in DP organoids suggesting that these PDOs may not retain the molecular profile of basal-like PDAC. LHSC124 again exhibited expression of both classical and basal-like genes consistent with its mixed morphology (**Figure 2C**).

Immunofluorescence (IF) microscopy for the basal-like specific marker, ANXA8, or the classical-specific marker CLDN18, revealed minimal ANXA8 expression in PDOs (**Figure 2D, Supplementary Figure 2**) and no ANXA8^+^ cells in GL PDOs, LHSC107 had an occasional ANXA8^+^ cell (<1% of cells), and LHSC124 had ANXA8^+^ cells appearing only in DP but not GL PDOs **(Figure 2D)**. Surprisingly, LHSC134 contained no ANXA8^+^ cells despite its DP morphology (**Supplementary Figure 2**). Conversely, all PDOs showed extensive CLDN18 expression **(Figures 2D** and **S2**), supporting transcriptomic analysis and suggesting components of the TME may be required to retain the basal-like molecular PDAC subtype.

### Analysis of Patient-Derived CAFs

The observation that LHSC39, which was established in monolayer co-cultures, had the most overt basal-like expression profile suggests non-cancer cells within the TME may support this molecular profile. Previous studies suggest CAF subtypes, such as iCAFs and myCAFs, promote PDAC subtypes.^25,26^ 2D cultures were examined for the presence of fibroblast subtypes initially through co-IF with the pan-CAF marker vimentin, or the myCAF marker, α-smooth muscle actin (αSMA; **Figure 3A**). Different expression patterns of αSMA were observed within and between samples, confirming fibroblast heterogeneity between patients. CAF57 and CAF62 showed minimal αSMA^+^ expression, while most cells in CAF10 and 73 cells showed prominent αSMA expression. To determine if αSMA-negative fibroblasts could be activated, CAF cultures were incubated with TGF-β, which promotes αSMA expression (**Supplementary Figure 3A**). An increase in αSMA^+^ cells was observed in all cultures (**Supplementary Figure 3B**) consistent with fibroblast activation. However, even after 24 or 48 hours of TGF-β treatment, many cells remained αSMA-negative, with the proportion of αSMA-negative cells ranging markedly between different patient samples.

**Figure 3.**
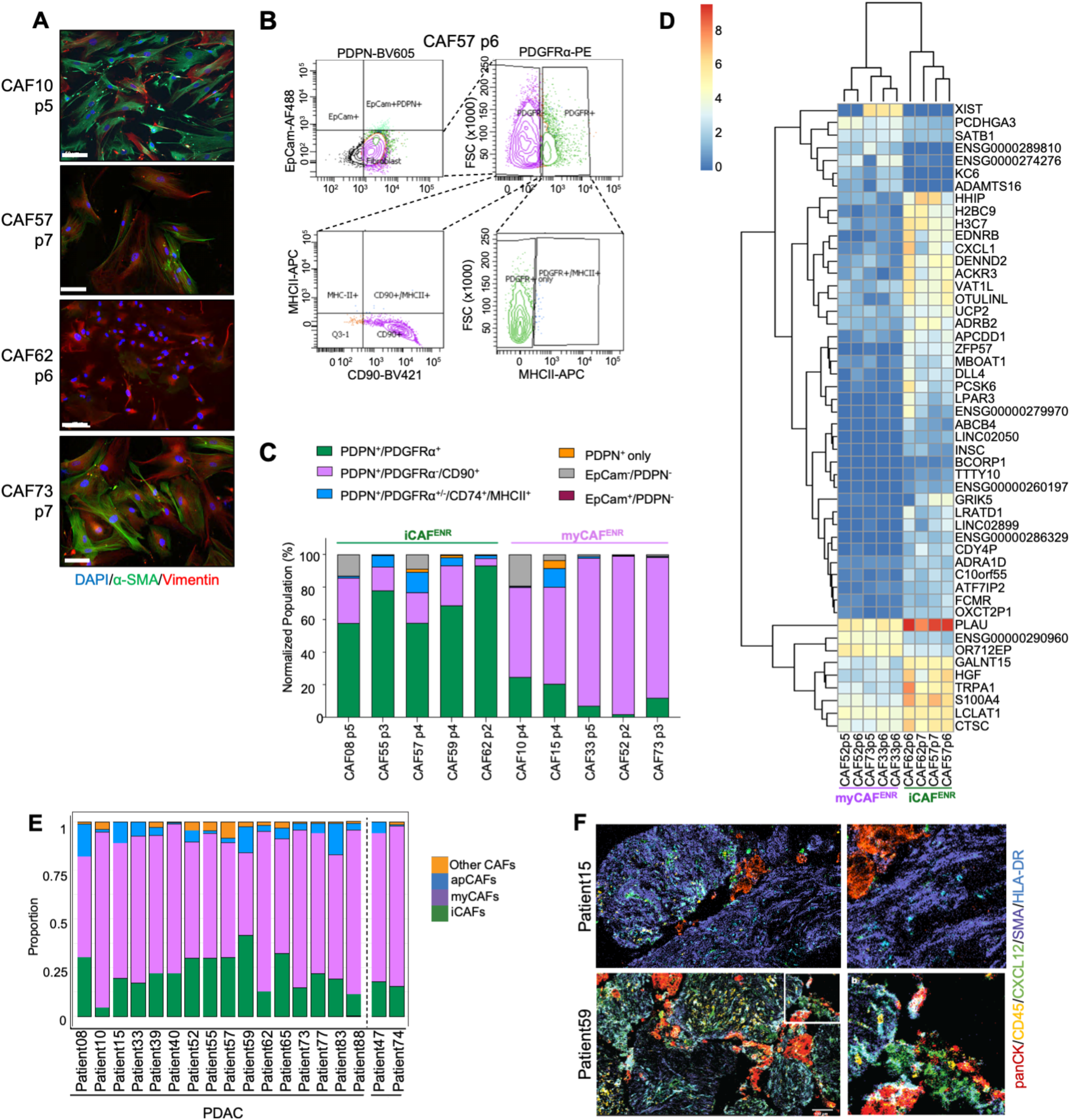
**(A)** Immunofluorescence staining of four primary CAFs. Vimentin, a panCAF marker is stained in red, α-SMA, a myCAF marker is stained in green, and nuclei are stained in blue with DAPI (scale bar = 100 µm). **(B)** Flow cytometry gating strategy in primary CAFs. First, CAFs are selected with PDPN^+^ staining. After CAF subpopulations are split by presence or absence of PDGFRα marker. Finally, cells are split with CD90^+^ and CD74^+^ or MHCII^+^ marker. **(C)** Synthesis of flow cytometry phenotyping on ten primary CAFs. PDPN^+^/PDGFRα^+^ represent iCAF subpopulation, PDPN^+^/PDGFRα^-^/CD90^+^ represent myCAF subpopulation and PDPN^+^/PDGFRα^+/-^/CD90^+/-^/CD74^+^/MHCII^+^ represent apCAF subpopulation. These results show two main groups of CAFs: iCAF enriched and myCAF enriched subpopulations. **(D)** Heatmap of top 50 genes differentially expressed between myCAF^ENR^ and iCAF^ENR^ subpopulations. iCAF^ENR^ subpopulations are clustering together and highly express genes like CXCL1 or PLAU. **(E)** IMC-derived per-patient cell composition proportions across 18 EUS samples. Bars show stromal CAF subtypes iCAFs, myCAFs, apCAFs, Other CAFs. Patients vary widely in total cellularity and in the relative mix of CAF populations. **(F)** Representative IMC images from two patients (15 and 59) illustrating spatial context. *Patient15*: contains stromal dense regions enriched in myCAFs (SMA-high) adjacent to tumor regions. *Patient59*: contains a sample enriched in iCAFs (CXCL12-high).

Next, flow cytometry was performed to identify the most common CAF subtypes. Staining for the pan-CAF marker, podoplanin (PDPN), was combined with myCAF (CD90), iCAF (PDGFRα) and apCAF (CD74 or MHCII) – specific markers. An antibody for EpCAM was included to identify cancer cells (**Figure 3B, C**). Analysis was performed on a range of passages from P2 to P7. Initial analysis of confirmed different proportions of CAF subtypes between patient samples consistent with IF analysis. Similar to IF analysis, ∼25% of cultures contain EpCAM+ cells (data not shown). The percentage of PDPN^+^ fibroblasts that were myCAFs (CD90^+^PDGFRα^-^) ranged from 4.5 to 97.4% while the proportion of iCAFs (CD90^-^PDGFRα^+^) ranged from 1.8 to 93.2%. In all cases, smaller proportions of apCAFs (CD74^+^ or MHCII^+^<5%) were observed along with PDNP^+^ cells not categorized into any CAF subtype. These findings are consistent with published single cells (sc) RNA-seq data suggesting CAF subtypes not categorized as myCAF, iCAF or apCAF, exist.^15,27,28^ To confirm these classifications, RNA-seq was performed on a subset of iCAF-enriched (CAF57 and 62, iCAF^ENR^) and my-CAF enriched (CAF33, 52 and 73, myCAF^ENR^) samples (**Figure 3D**). As predicted by flow analysis, myCAF^ENR^ and iCAF^ENR^ samples clustered away from each other and iCAF-related genes, including *PLAU* and *CXCL1*, were higher in the iCAF^ENR^ samples.

Next, to assess whether CAF cultures retained patient-specific *in vivo* characteristics, imaging mass cytometry (IMC) was performed on tissue sections from the same patient biopsies used to derive the cultures (**Figure 3E, F; Supplementary Figure 3C-E**). Sixteen patient samples were assessed alongside two pancreatitis samples (CAF47 and CAF74) as controls. Ten of these cases corresponded to CAF cultures classified by flow cytometry (**Figure 3C**), allowing for matched comparison. The heterogeneity observed across CAF cultures by flow cytometry and IF was similarly evident to that observed in the matched patient tissue by IMC. To directly compare with the *ex vivo* CAF classifications, iCAF abundance was quantified as a proportion of total CAFs in each patient sample. CAF10 and CAF62 were excluded from quantitative IMC comparisons due to low total segmented cell counts which limited cellular representation for estimating CAF subtype proportions. Although myCAFs remained the predominant CAF population across samples, specimens classified as iCAF^ENR^ in culture (CAF08, 55, 57, and 59) showed higher relative iCAF representation, ranging from 29.9–41.5% of the total CAF population, compared with 14.8–29.8% in samples identified as myCAF^ENR^ in culture (CAF15, 33, 52 and 73) (**Figure 3E**; exact Mann-Whitney U test, p<0.03). These findings suggest that the relative iCAF enrichment observed *ex vivo* is reflected in matched patient tissue, despite the overall predominance of myCAFs *in situ*.

### CAF-conditioned media alters PDO phenotypes

The observation that PDOs retained morphologies similar to their *in vivo* phenotypes but showed loss of basal-like specific markers suggests components of the TME affect the molecular profiles of cancer cells. Therefore, we examined whether conditioned media (CM) from the various patient-derived CAF cultures could promote changes in PDO phenotype. Early passage CAFs samples (P2-P7) were grown to 80% confluency to generate CM. Samples with no tumor cells present were chosen, and four iCAF^ENR^ or four myCAF^ENR^ samples used. CM was filtered to ensure no cells were inadvertently transferred and combined at a 1:1 ratio with normal organoid media.

LHSC04 (GL) and LHSC107 (DP) PDOs were incubated with CM or with non-conditioned CAF media mixed with PDO media. PDOs were followed for two weeks and assessed for growth and morphology (**Supplementary Figure 4B**). Two weeks after treatment, changes were observed in LHSC04 growth and morphology (**Figure 4A**). iCAF^ENR^-CM promoted denser PDO morphology (**Figure 4A**) with histological analysis confirming a shift from simple cuboidal epithelium to a columnar or even stratified epithelium (**Figure 4A, B**). CM from myCAF^ENR^ samples showed no effect on GL PDO morphology but did increase the size of GL PDOs (**Figure 4A-C**). No significant changes in morphology were observed in DP organoids (**Supplementary Figure 4C-E**). Analysis of other GL PDO lines, LHSC137 and LHSC143, resulted in similar outcomes, with iCAF^ENR^-CM promoting a more stratified or dense appearance (**Figure 4 D-F**), while LHSC124 and LHSC134 showed no morphological differences (**Supplementary Figure 4 C-E**). Analysis of KI67 expression showed no consistent increases in PDO proliferation based on treatment. In LHSC04 PDOs, Ki67 was only significantly different between myCAF^ENR^ and iCAF^ENR^ treatments (**Supplementary Figure 5**). In LHSC107, the number of Ki67^+^ cells was increased only following myCAF^ENR^ treatment (**Supplementary Figure 5**).

**Figure 4.**
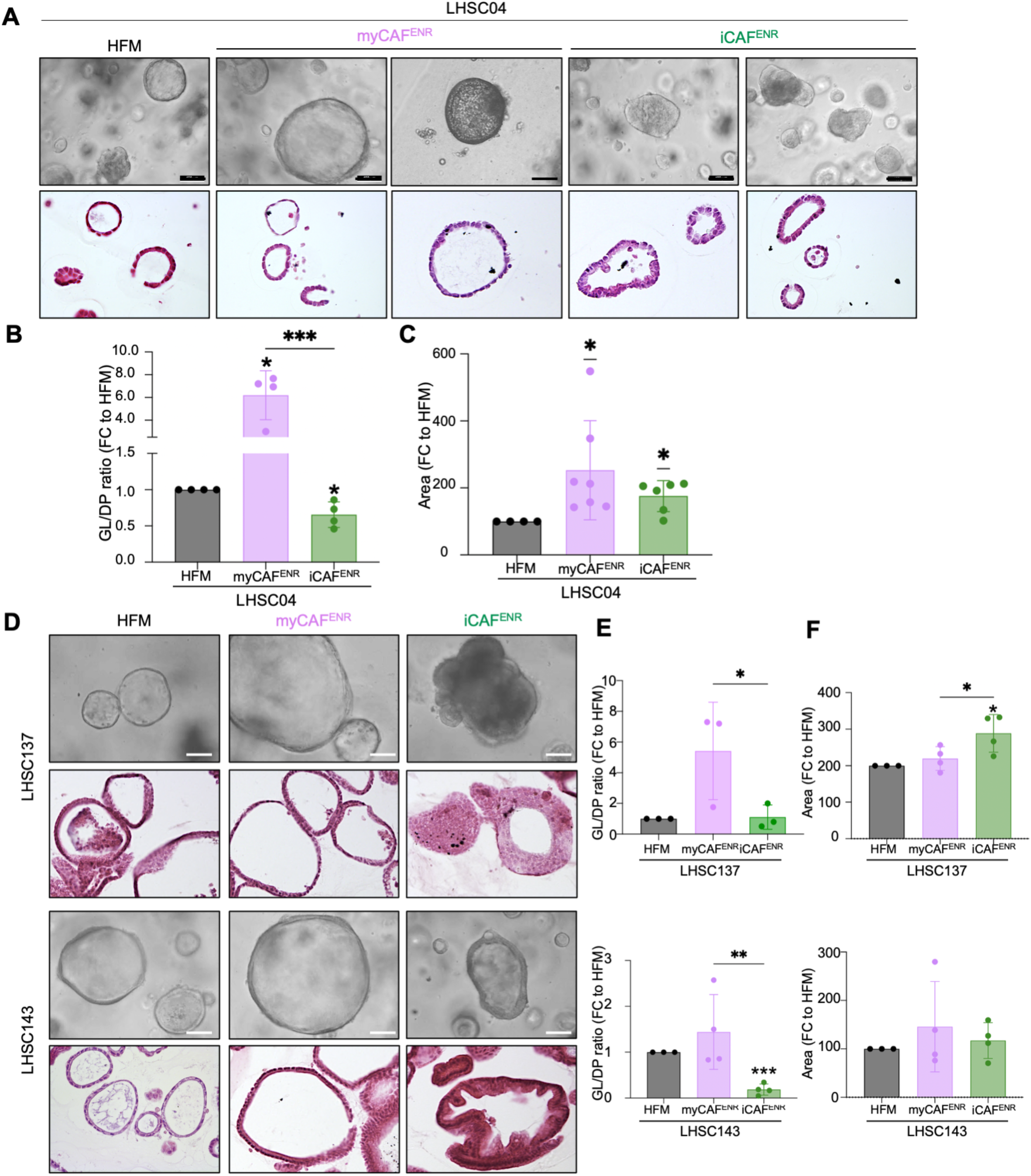
**(A)** LHSC04 PDO brightfield and H&E (Scale bar = 100µm) untreated (HFM) or treated two weeks with iCAF^ENR^ or myCAF^ENR^ conditioned media (CM). iCAF^ENR^-CM induces morphological changes in LHSC04. **(B)** Quantification of GL/DP ratio on LHSC04 PDO after two weeks of treatment with iCAF^ENR^ or myCAF^ENR^-CM. **(C)** Quantification of area ratio on LHSC04 PDO after two weeks of treatment with iCAF^ENR^ or myCAF^ENR^-CM. **(D)** LHSC137 and 143 PDOs brightfield and H&E (Scale bar = 100µm) untreated (HFM) or treated two weeks with iCAF^ENR^ or myCAF^ENR^ conditioned media (CM). iCAF^ENR^-CM induces morphological changes in both PDOs. **(E)** Quantification of GL/DP ratio on LHSC137 and 143 PDOs after two weeks of treatment with iCAF^ENR^ or myCAF^ENR^-CM. **(F)** Quantification of organoid area ratio on LHSC137 and 143 PDOs after two weeks of treatment with iCAF^ENR^ or myCAF^ENR^-CM. *p≤0.05, **p≤0.01, ***p≤0.001.

Since GL-PDOs showed changes in epithelial morphology after treatment with iCAF^ENR^-CM consistent with a more DP phenotype, the expression of basal-like markers, KRT17 and ANXA8, were compared by immunofluorescent analysis (**Figure 5**). Treatment with iCAF^ENR^-CM but not myCAF^ENR^-CM produced an increase in KRT17 (**Figure 5A**) and ANXA8 (**Figure 5B**) accumulation in both GL and DP organoids relative to media treatment alone. Conversely, myCAF^ENR^-CM promoted minimal changes in the expression of either KRT17 or ANXA8 accumulation, except for increased KRT17 accumulation specifically in LHSC107. Similar analysis of the mixed LHSC124 organoids and the GL LHSC137 and 143 showed increase expression of KRT17 and ANXA8 after iCAF^ENR^ CM **(Supplementary Figure 6A and 6B**).

**Figure 5.**
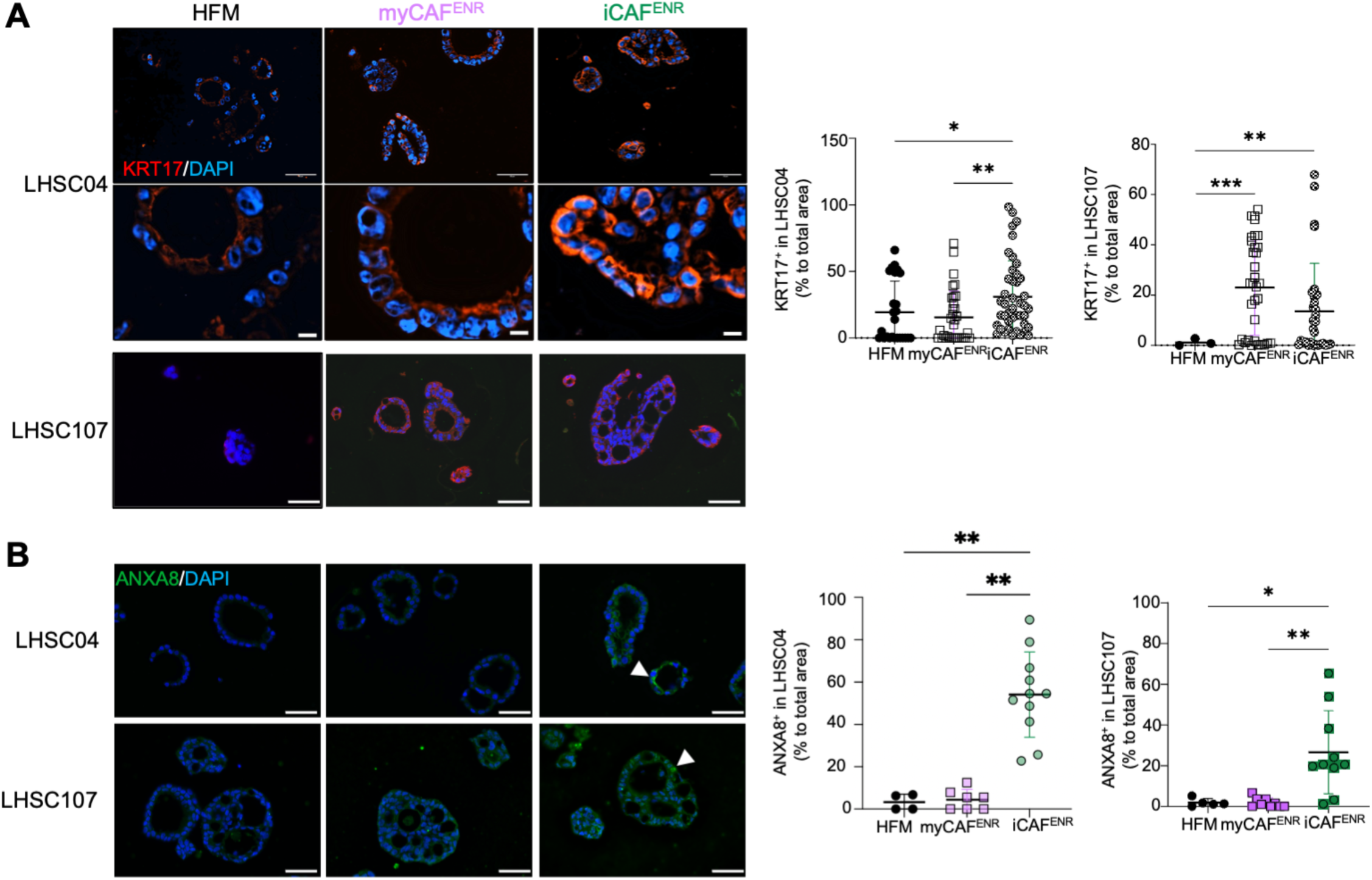
Immunofluorescence staining of **(A)** KRT17 in red or **(B)** ANXA8 in green and of nuclei in blue by DAPI in two different PDOs untreated (HFM) or treated for two weeks with iCAF^ENR^ or myCAF^ENR^-CM. KRT17 and ANXA8 are basal markers and show increased staining in iCAF^ENR^-CM treated PDOs (scale bar = 50 µm). *p≤0.05 **p≤0.01 ***p≤0.001.

### iCAF^ENR^-CM alters the molecular landscape of GL-PDOs

To investigate the molecular consequences by which iCAF^ENR^-CM promotes a basal-like phenotype, RNA-Seq was performed following treatment of PDOs (**Supplementary Figures 7A, Figure 6A**). Gene expression was compared between two DP PDOs (LHSC107, 134) and three GL PDOs (LHSC04, 137, 143) two weeks into treatment with conditioned media from two myCAF^ENR^ (CAF33, CAF52) or iCAF^ENR^ (CAF57, CAF62). As a control, RNA-seq was performed on PDOs treated with CAF/PDO media but not conditioned by CAFs (**Supplementary Figure 7A**). Changes in gene expression in DP PDOs were modest following exposure to CM from either myCAF^ENR^ (164 differentially expressed genes (DEGs); logFC >1, p<0.05) or iCAF^ENR^ (204 DEGs) samples (**Supplementary Figure 7B**). Similarly, negligible differences were observed comparing GL PDOs treated with CM from myCAF^ENR^ samples (**Figure 6A**). Conversely, PDOs treated with CM from iCAF^ENR^ samples resulted in marked changes in gene expression (643 DEGs decreased and 362 DEGs increased) compared to non-conditioned media (**Figure 6A**). REACTOME analysis using these DEGs identified enrichment of gene repression pathways including histone deacetylases (HDACs) and DNA methylation, and chromatin/chromatid regulators (**Supplementary Figure 7C**) which suggest epigenetic repression may account for the higher number of genes repressed by iCAF^ENR^-CM. To examine the effect of iCAF^ENR^-CM treatment on chromatin accessibility, we performed ATAC-seq (**Figure 6B, C**) at the same time points as transcriptomic analysis. Global decreases in chromatin accessibility were observed in PDOs treated with iCAF^ENR^-CM (**Figure 6B**). Targeted examination of genomic regions surrounding classical or basal-like genes revealed chromatin accessibility patterns indicative of a classical phenotype in GL PDOs LHSC04 and LHSC137, with classical genes (i.e. CLDN18, GATA6 and TFFs) showing open chromatin and basal-like genes (i.e ANXA8 and KRT5) having closed chromatin. Treatment with iCAF^ENR^ CM decreased chromatin accessibility at classical genes suggesting epigenetic mechanisms may restrict expression of these genes. Surprisingly, the TSS regions at basal genes showed more variable accessibility after the same treatment (**Figure 6C**).

**Figure 6.**
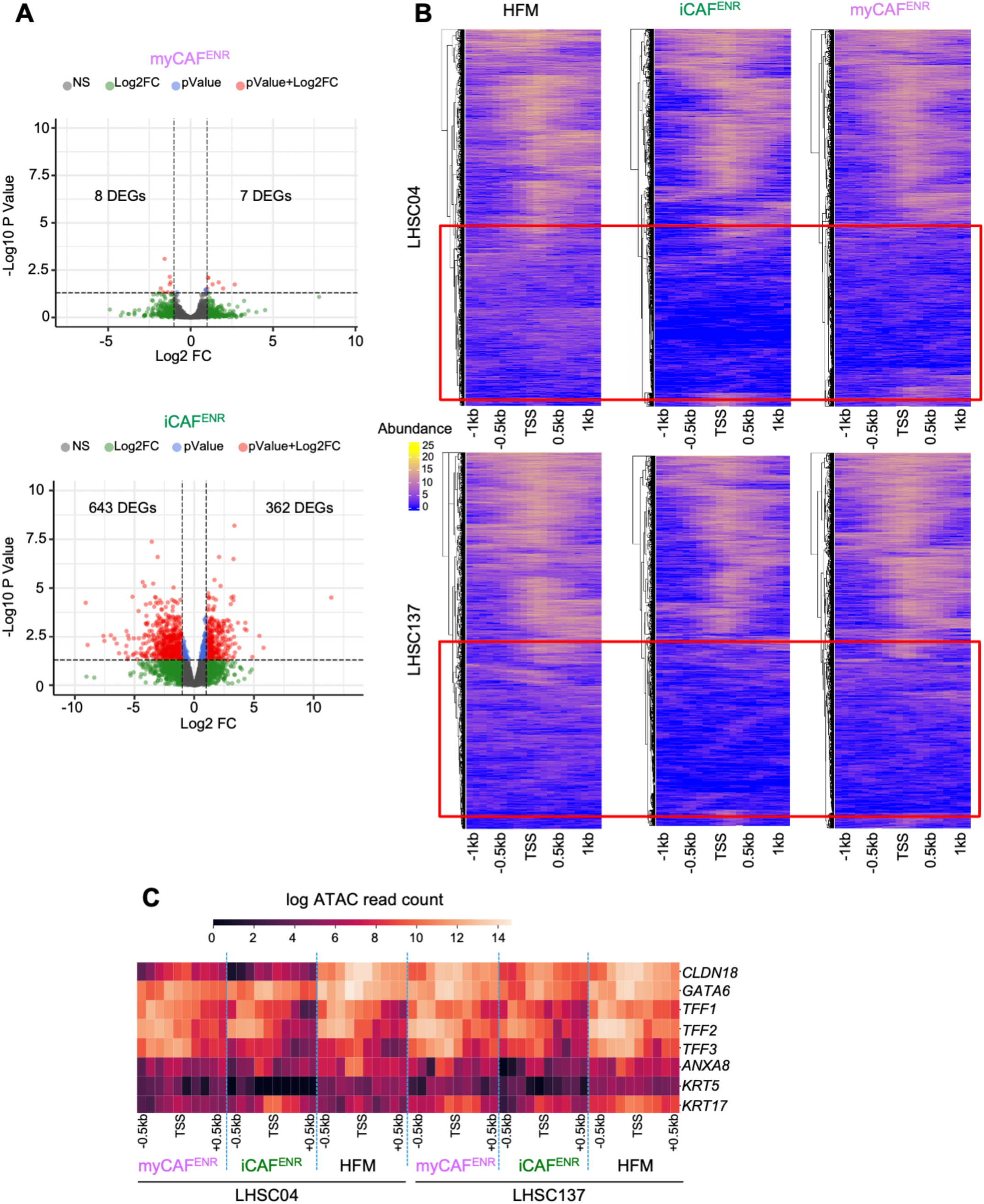
**(A)** Volcano Plot of differentially expressed genes in GL-PDOs (LHSC04, 137 and 143) treated for two weeks with two different iCAF^ENR^ or myCAF^ENR^-CM. iCAF^ENR^-CM treated GL-PDOs show a high downregulation of gene expressions (643 genes). **(B)** ATAC Sequencing analysis of (top) LHSC04 and (bottom) LHSC137 PDOs untreated (HFM) or treated two weeks with iCAF^ENR^ or myCAF^ENR^-CM. iCAF^ENR^-CM increases closing regions (blue color) compared to both HFM and myCAF^ENR^-CM (red square). **(C)** ATAC-Sequencing read counts from −0.5 to 0.5 kb from TSS of classical and basal genes in LHSC04 and LHSC137 treated for 2 weeks with with iCAF^ENR^ or myCAF^ENR^-CM.

### Effects of iCAF^ENR^-CM in GL-PDOs are linked to metabolites production

The levels of IL-6 and TGF-β, which promote different CAF cancer cell phenotypes^29–32^, were assessed by ELISA (**Supplementary Figure 8A**) and showed no significant differences between iCAF^ENR^ and myCAF^ENR^ media. Therefore, to identify iCAF^ENR^-derived factors promoting morphological changes in GL-PDOs, metabolomic analysis was performed on CAF-CM (**Figure 7**). Eight CAF samples were assessed (**Figure 7A**) and nsupervised analysis separated these samples into iCAF^ENR^ CAF (CAF08, 57, 59 and 62) and four myCAF^ENR^ CAF (CAF10, 33, 52 and 73) groups. The top 25 most significantly different metabolites identified taurine (2-fold +/-) and glyceraldehyde-3-Phosphate (G3P; X-fold +/-) as enriched in iCAF^ENR^ CM while taurocholic acid was significantly increased in myCAF^ENR^ CM (**Figure 7B, C, Supplementary Figure 8 B, C**). KEGG pathway analysis using this metabolomic data confirmed significant increases in taurine and hypotaurine metabolism specifically in iCAF^ENR^-CM compared to myCAF^ENR^-CM (**Figure 7D, Supplementary Figure 8D**) and metabolomic analysis on intracellular components of iCAF^ENR^ and myCAF^ENR^ samples also confirmed increased taurine production in iCAF^ENR^ samples linked to hyperactive glycolysis and the pentose phosphate pathway which would increase NADPH generation, which is essential for production of taurine dioxygenases (**Supplementary Figure 8E**). Combined, this analysis suggests that iCAF^ENR^ CAF samples produce metabolites to promote a basal-like phenotype in pancreatic cancer PDOs.

**Figure 7.**
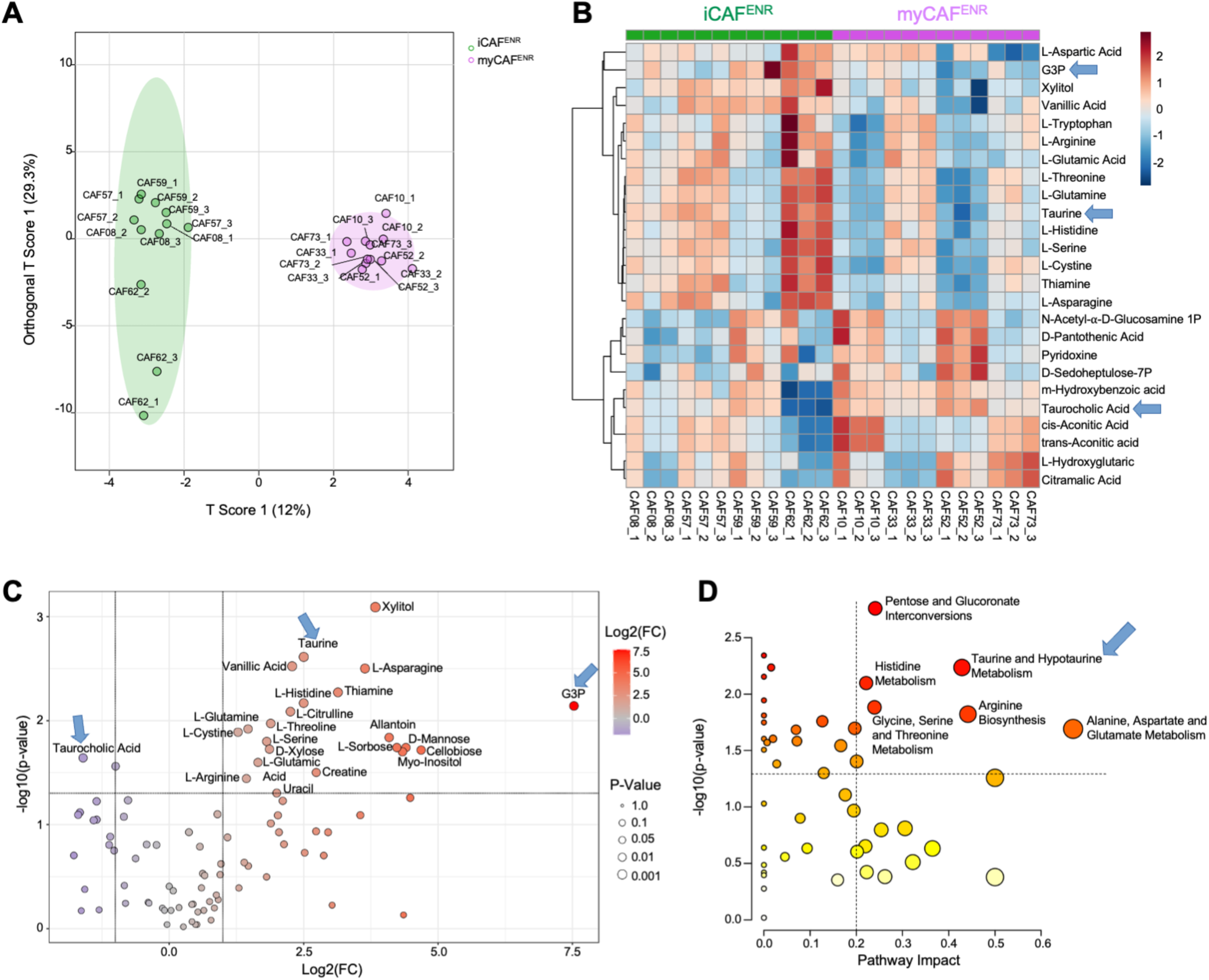
**(A)** Orthogonal PLS-DA analysis of metabolomic data on four iCAF^ENR^ and four myCAF^ENR^ primary supernatants (in triplicate). The results show a strong separation between iCAF^ENR^ and myCAF^ENR^ conditioned media. **(B)** Top 25 metabolites dysregulated between iCAF^ENR^ and myCAF^ENR^ primary supernatants. These results identify G3P and Taurine as overexpressed in iCAF^ENR^ vs myCAF^ENR^ samples and Taurocholic Acid as downregulated in iCAF^ENR^ vs myCAF^ENR^. **(C)** Volcano Plot of differentially expressed metabolites in iCAF^ENR^ vs myCAF^ENR^ supernatants (FC≥2, p-value≤0.05). Taurine and G3P are showing as significantly enriched in iCAF^ENR^ supernatant while Taurocholic Acid is significantly decrease in iCAF^ENR^ supernatant. **(D)** Enriched KEGG pathway analysis in iCAF^ENR^ vs myCAF^ENR^ supernatants (lmpact≥0.2, p-value≤0.05). Taurine and Hypotaurine metabolism are significantly increased in iCAF^ENR^ conditioned media.

## Discussion

Pancreatic ductal adenocarcinoma is a devastating disease due to late diagnosis, rapid progression and resistance to therapy. One major issue in treating PDAC is the heterogeneity observed within and between patients. Multiple subtypes based on transcriptional and phenotype profiles exist for both cancer cells and non-cancer cells in the tumor microenvironment (TME). While patient-specific treatment is needed in PDAC, there are limited opportunities to match patients to therapy. Recent successes in developing patient-derived organoids (PDOs) have provided some hope in personalized medicine^33–35^, but this approach requires obtaining tumors in a timely fashion. In addition, studies have been mixed regarding translation of *ex vivo* findings to *in vivo* biology. To this end, we developed a program that obtains tissue at the time of diagnostic endoscopic ultrasound (EUS) and includes derivation of cancer and non-cancer cells under 3D and 2D conditions. Our analysis suggests 3D conditions favor a classical phenotype based on molecular profiling, and that the presence of CAFs affects this profile. Based on this work, we suggest translation of PDO sensitivities to treatment should consider the effects of CAFs on cancer cells.

The use of patient-derived organoids (PDOs) is gaining credibility for understanding specific therapeutic vulnerabilities in cancer. Using PDOs for PDAC has been problematic due to the paucity of patients eligible for surgery, a major source for cells used to develop PDOs and the rapid progression of this cancer. We have addressed these issues by establishing PDOs from EUS at the time of initial diagnosis. EUS captures all PDAC patients prior to treatment, unlike surgery in which most patients have received neoadjuvant therapy, which may negatively affect the ability to develop PDO lines. Over the course of this study, we reported a success rate of ∼12.3% for developing PDO lines from EUS samples derived from PDAC. While at the lower end of published success rate, several factors contribute to this rate of success. PDOs developed from EUS have been described in several studies, with successes ranging from 34 up to 84%^36–39^. However, these studies were based on limited numbers and did not indicate whether PDO lines could be established over multiple passages. Non-cancerous tissue from EUS can be cultured effectively and propagated dependent on the culture media used. Our initial success at P1 is >80% and may reflects other components of the biopsy (i.e. intestinal epithelium). In culture media that specifically promotes cancer cell growth, PDOs will not be maintained to later passages (i.e. past P5). A recent study described high success rates (70%) at earlier passage, which decreased to as those samples reached passage 5^37^, while a second study showed a 34% success rate for PDOs reaching 5 passages.^36^ In addition, PDOs in previous studies were often validated by histology only (H&E staining) with no molecular characterization, including mutational status presented. We provide molecular profiling that confirms a range of subtypes and genetic mutations consistent with the *in vivo* phenotype. All PDOs were validated multiple methods including mutation identification by Sanger sequencing or Oncomine sequencing, histological analysis (H&E and immunofluorescence staining) and RNA-seq analysis. This level of validation ensured all lines represent true PDAC PDOs.

Another factor that may account for lower success rates is the technical aspects of EUS guided tissue acquisition and organoid cultures. A critical component to success will require consistency in EUS sampling and organoid isolation and culturing. A prospective study examining EUS procedural factors affecting organoid growth is currently underway. Of the 202 samples provided, 61 samples (30.2%) were discarded for various technical reasons. Additionally, while our protocols followed published standards, optimization of isolation and culture conditions was required based on informal discussions with other centers developing organoid lines. We now provide the most effective approach observed in our center with the highest success rates over the last 40 attempts (>30%). Based on our experience, it is imperative isolation protocols and cell culture conditions be precisely described and extensive PDOs characterization performed to confirm the presence of tumor cells.

Cancer-associated fibroblasts represent between 20 to 50% of all non-cancer cells in PDAC tumor^27,40^ and several studies showed CAFs impact cancer cell growth, gene expression and resistance. In most cases, these studies focused on individual CAF subtypes or did not perform characterization of the patient CAFs used. EUS sampling allowed us to develop heterogeneous, patient-derived cancer-associated fibroblasts that could be matched to PDOs. Similar to PDOs, CAFs retained *in vivo* phenotypes for several passages regarding the relative amount of iCAFs and myCAFs present in the tissue. While iCAFs and myCAFs were the predominant CAF populations, antigen-presenting CAFs and CAFs not belonging to these subsets (i.e. PDPN^+^ only) were also observed. Recent scRNA-seq identified additional CAF subtypes, which may account for this population, as well as further sub-divide the groups we observed.^15,27,41,42^

Using these heterogeneous CAF samples, we showed conditioned media from iCAF^ENR^ cultures promoted a basal-like phenotype within PDOs. Morphologically, myCAF^ENR^ and iCAF^ENR^-CM showed no overt effects on basal PDOs, and changes to the transcriptomes of GL PDOs after iCAF^ENR^-CM treatment were muted in DP PDOs. These findings suggest differences in PDO sensitivities reflect morphology and may be tied to epigenetic programming within the cells. These findings are reflected by iCAF^ENR^-CM significantly increasing basal marker expression of KRT17^43,44^ or ANXA8^45^ in both DP and GL PDOs despite having a more limited effect on the molecular profile of DP PDOs. Recent studies suggest iCAFs are linked to a classical phenotype^26^, which is not consistent with our findings. In that study, conditioned media and co-culture of CAFs promoted a basal-like shift in patient matched lines of CAFs and cancer cells. The authors interpreted this effect to myCAFs, but the CAFs and cancer cells used in the study were not fully characterized for potential heterogeneity and confirmation of high myCAF percentages within the cultures was not provided. Conversely, analysis of 2D pancreatic cell lines showed iCAF secretion enhanced colony formation and invasion abilities, two features associated with basal-like subtype.^46^ iCAFs have immunosuppressive effects on the TME through cytokine secretion of IL-6 or CXCL12^47,48^ and induce shifts to glycolytic metabolism in cancer cells.^12,49^ These findings suggest iCAFs support PDAC aggressiveness, consistent with a more basal-like phenotype. It is possible that the CAF populations promoting PDO reprogramming represent a unique population or the combined effects of multiple cell types. Future experiments should use scRNA-seq analysis to define specific CAF populations promoting these changes.

### iCAF^ENR^ lines drive the basal-like subtypes through metabolic signaling and epigenetic modulation

Initial analysis of the molecular mechanisms driving iCAF^ENR^ CM changes into PDOs suggest secreted metabolites are responsible morphological changes observed in classical organoids. We identified iCAF^ENR^ cultures were taking up less amino acids than myCAF^ENR^ cultures suggesting distinct metabolic dependencies between populations, correlating with previous findings.^40^ Increases in glyceraldehyde-3-phosphate (G3P) in iCAF^ENR^-CM, which would up-regulate glycolytic flux^50^, and taurine, which has been linked to increased PDAC progression, were observed in iCAF^ENR^ cultures. Taurocholic acid (TCA), which is formed by taurine and cholic Acid, significantly decreased in iCAF^ENR^-CM. These results suggest a population of cells in the iCAF^ENR^ cultures prioritize unconjugated taurine over TCA in PDAC cells. This switch would favor cysteine/glutathione support, as taurine is part of the transsulfuration pathway^51^, and suggest iCAF^ENR^ populations promote redox/glycolysis pathways in PDAC cancer cells.

Our metabolomic analyses support a model where iCAF^ENR^ cells have active glycolysis and PPP pathways, which would lead to production of NADPH and GSH, and enter the transsulfuration pathway to favor taurine export. Glycolysis and PPP pathways were hyperactive, with specific increases in dihydroxyacetone phosphate, D-fructose-6P, phosphoenolpyruvic acid, arabinose-5P and 2-deoxyguanosine-5P. Such a switch would increase NADPH generation, which is essential for GSH production and taurine dioxygenases.^52^ GSH is overproduced in iCAF^ENR^ samples, reinforcing a model in which the transsulfuration pathway plays an essential role in iCAF^ENR^ metabolism. Increases in L-citrulline and 2-ketobutyrate further support transsulfuration pathway activity. Previous work by Zhu et al., showed increased GSH favored dual export of cysteine/taurine to abort tumor ferroptosis, supporting the taurine production observed in iCAF^ENR^-CM.^51^

Finally, this work suggests that adding CAFs back to PDOs promotes epigenetic reprogramming that favors gene repression. Previous studies also show extracellular taurine enhances GSH production in PDAC through the transsulfuration pathway, thereby reducing ROS and enhancing demethylation of active histone marks. This would result in reduced DNA accessibility, which is suggested by our ATAC-Sequencing data.^53,54^ Interestingly, conditioned media from all CAF samples show decreased chromatin accessibility. We are currently examining whether long term culturing of PDOs leads to de-repression at the epigenetic level, and whether this alters therapeutic responses over time.

To conclude, we provide methodology and an experimental framework needed to rapidly characterize PDAC tumors from patients to provide translational potential for treatment. We show that PDOs reflect in vivo characteristics but that the TME, and specifically CAFs, need to be considered when understanding patient-specific heterogeneity and biology. We also show potential links between metabolic changes and epigenetic reprogramming that dictate PDO subtypes and affected by the presence of CAFs.

## Funding and Acknowledgements

This work would not have been possible without financial support from the Canadian Institutes of Health Research (MOP#PJT166029), the Cancer Research Society of Canada and the Rob Lutterman Foundation for Pancreatic Cancer Research (no grant number), the Ontario Institute of Cancer Research, and the Canadian Cancer Society. We are especially indebted to support from the Baker Centre for Pancreatic Cancer. EJP and FB was supported by Mitacs Accelerate Fellowships. MN was supported by an Ontario Graduate Scholarship. The funder played no role in study design, data collection, analysis and interpretation of data, or the writing of this manuscript. Finally, the authors would like to thank the patients who donated their tumour biopsies for this work.

## Author Contributions

EJP - data acquisition and interpretation, manuscript writing and editing; FAB - data interpretation, manuscript writing and editing; MN - data acquisition; CB - data acquisition; JC - data acquisition; JG - data acquisition and manuscript writing; NS - data acquisition and manuscript writing; NA – data acquisition; PJ - data acquisition; LTPK - data acquisition; PS - data acquisition and manuscript writing; GG – data acquisition; DD – data acquisition; LJM – data acquisition; DO – data acquisition; CL - - funding support, data acquisition supervision and manuscript editing; MC – data acquisition and supervision, editing manuscript; KL – funding support and data acquisition; SW – funding support and data acquisition; HJ - funding support, data acquisition supervision and manuscript editing; NH – patient access and sample acquisition; MS - patient access, sample acquisition and manuscript editing; BY - patient access, sample acquisition and manuscript editing; CLP – study design funding support, data analysis and interpretation, supervision, manuscript writing and editing.

## Data Availability

All NGS data will be made available upon request. Raw data will be uploaded to NCBI upon acceptance.

