## Supplemental Figures for "Generation of tumor organoids and cancer-associated fibroblasts from minimal pancreatic cancer biopsies enables investigation of tumor-stroma interactions"

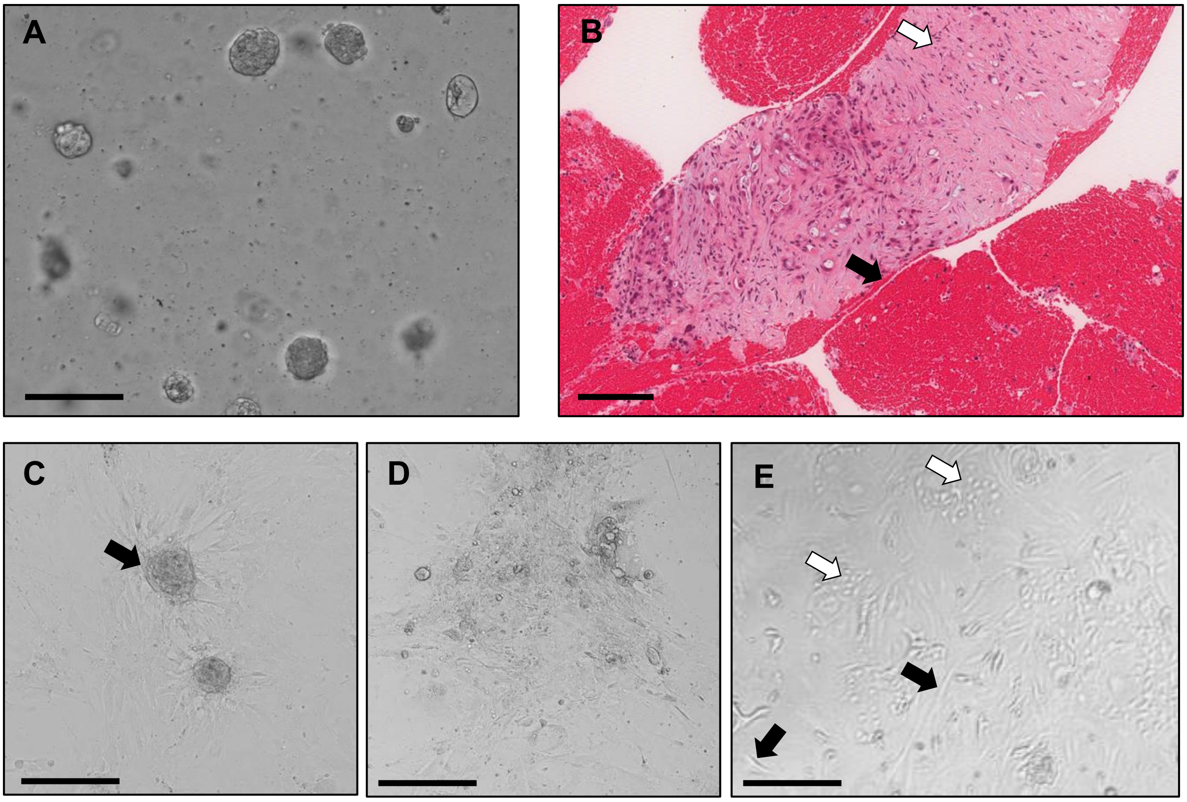


**Supplementary Figure 1 (Jaune-Pons et al. 2026). Characterization of primary cultures from endoscopy. (A)** Phase contrast images showing organoids at Passage 1. **(B)** FNB showing limited cancer cells in a fibrous background (white arrow) with blood cell accumulation (black arrow). Phase contrast of 2D cultures at P2 showing **(C)** putative organoids (black arrow) and **(D)** culture heterogeneity. **(E)** LHSC39 at P5 with putative fibroblasts (white arrows) and cancer cells (black arrows. Scale bar = 100 µM.

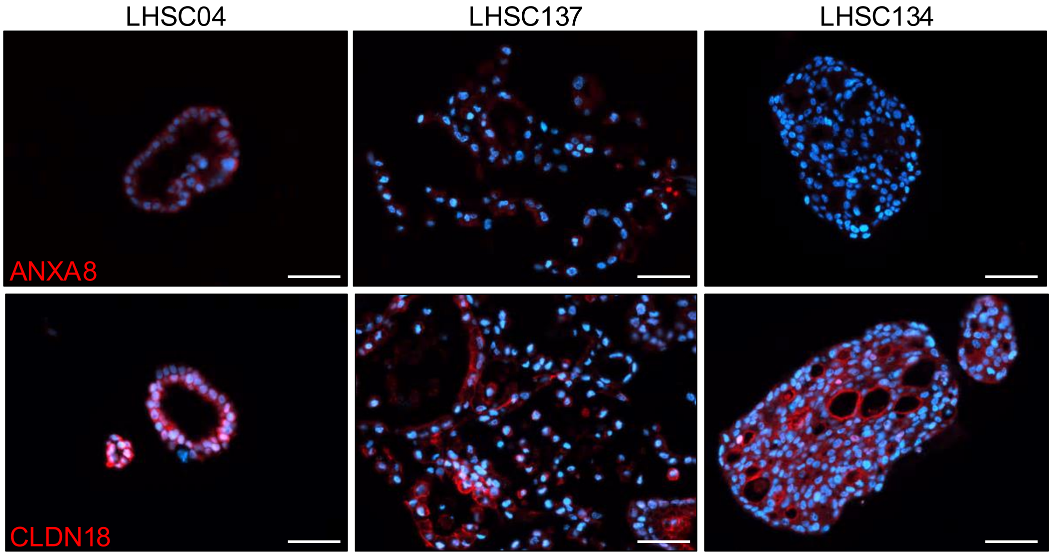


**Supplementary Figure 2 (Jaune-Pons et al.,2026). Expression of basal-like and classical markers in PDOs.** Immunofluorescence staining for the basal-like marker, ANXA8 (red; top row) or the classical marker, CLDN18 (red; bottom row) combined with DAPI staining of nuclei in three different organoids. Scale bar =50 µM.

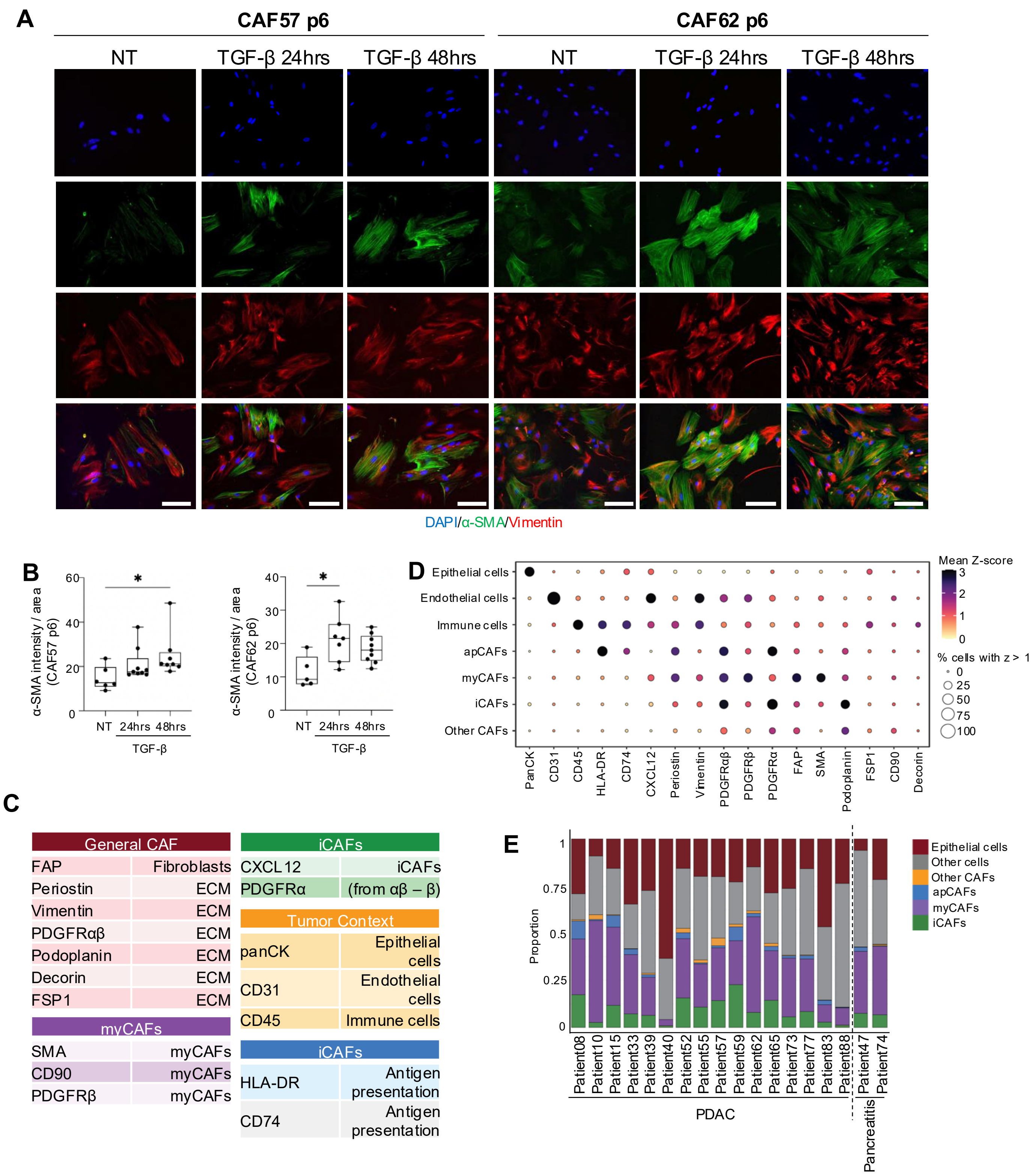


**Supplementary Figure 3 (Jaune-Pons et al., 2026). Characterization of cancer-associated fibroblasts in culture or in fine needle biopsies from endoscopic ultrasound. (A)** Immunofluorescence staining for vimentin (all CAFs, red) and α-SMA (myCAFs; green) in primary CAFs treated with 100 pM of TGF-β for 24 or 48 hours. Scale bar = 100 µm. **(B)** Quantification of α-SMA intensity before and after treatment with TGF-β for 24 or 48 hours. *p≤0.05 **(C)** IMC antibody panel and marker set used to define cell types and CAF subtypes in this study. **(D)** Bubble plot of marker expression by cell type. Dot color encodes mean standardized intensity (mean z-score); dot size represents the percentage of cells with z > 1. iCAFs are enriched for CXCL12/PDGFRα, myCAFs for SMA/CD90/PDGFRβ, apCAFs for HLA-DR/CD74; panCK, CD31, and CD45 track epithelial, endothelial, and immune compartments, respectively. **(E)** IMC-derived per-patient cell composition proportions across 18 EUS samples

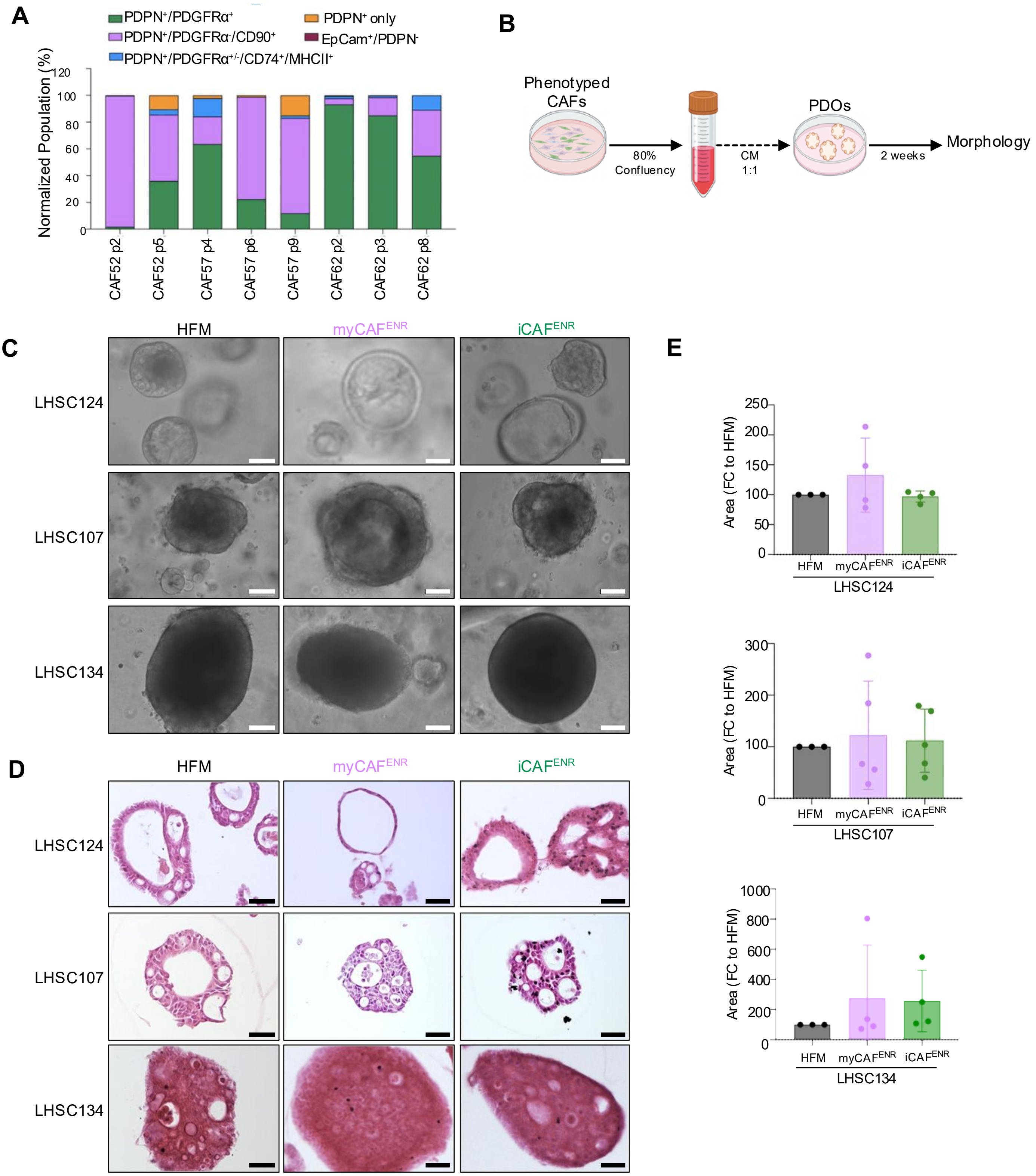


**Supplementary Figure 4 (Jaune-Pons et al., 2026). Characterization of PDOs following treatment with CAF-conditioned media. (A)** Flow cytometry phenotyping on primary CAFs at different passages (P) to show shifts in heterogeneity. PDPN$^{+}$/PDGFRα$^{+}$ represent iCAF subpopulation (green), PDPN$^{+}$/PDGFRα$^{-}$/CD90$^{+}$ represent myCAF subpopulation (pink) and PDPN$^{+}$/PDGFRα$^{+/-}$/CD90$^{+/-}$/CD74$^{+}$/MHCII$^{+}$ represent apCAF subpopulation (blue). Orange depicts PDPN+ cells that do not stain for any other CAF marker. **(B)** Scheme showing CAF-CM treatment of PDOs. Brightfield **(C)** or H&E **(D)** imaging of PDOs LHSC124, LHSC107 and LHSC134 (Scale bar = 100 µm) following incubation in human feeding media (HFM; control) or following treatment for two weeks with iCAF$^{ENR}$ or myCAF$^{ENR}$ conditioned media (CM). **(E)** Quantification of area ratio on LHSC124, 107 and 134 PDOs two weeks after treatment with iCAF$^{ENR}$ or myCAF$^{ENR}$-CM.

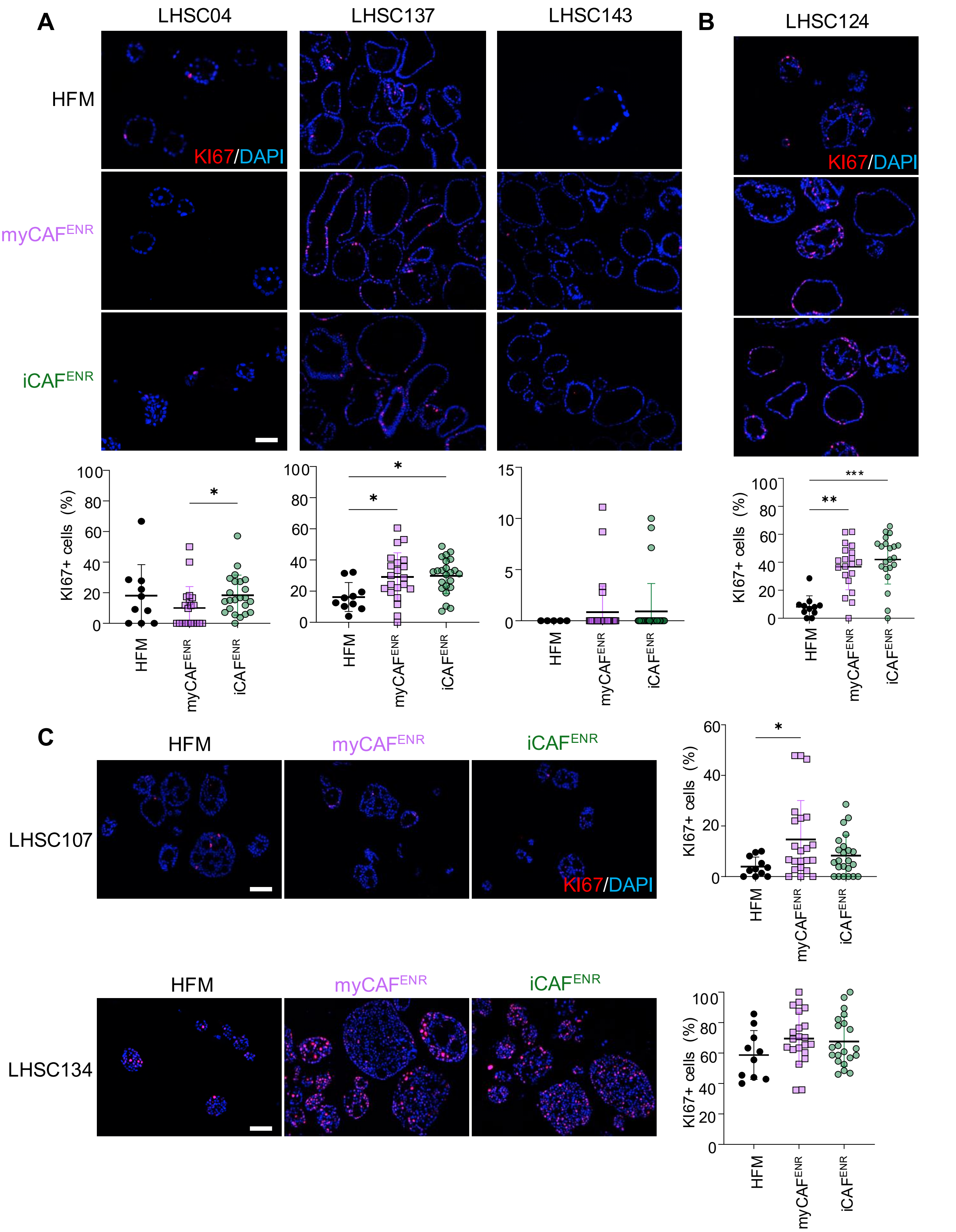


**Supplementary Figure 5 (Jaune-Pons et al., 2026). Analysis of PDO proliferation following treatment with CAF-conditioned media.** Immunofluorescence staining for KI67 (red) counterstained with DAPI (blue) in **(A)** GL-PDOs, **(A)** mixed PDOs, and **(C)** DP-PDOs treated for two weeks with human feeding media (HFM), iCAF$^{ENR}$,or myCAF$^{ENR}$-CM. Scale bar = 50 µm. *p≤0.05,**p≤0.01,***p≤0.001.

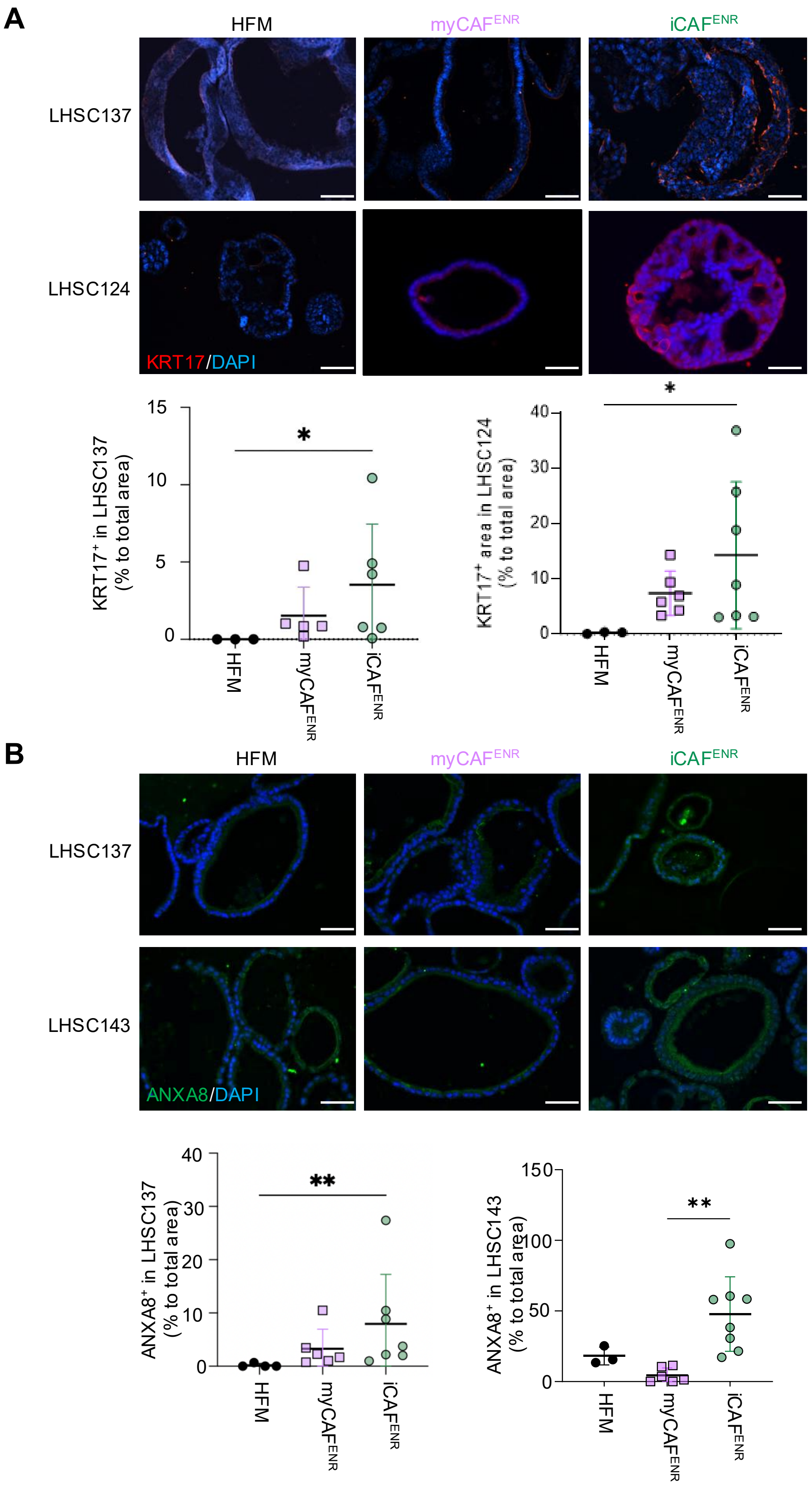


**Supplementary Figure 6 (Jaune-Pons et al.,2026). Analysis of molecular subtypes in PDAC PDOs.** Immunofluorescence staining for **(A)** KRT17 (red) or **(B)** ANXA8 (green) counterstained with DAPI (blue) in PDOs grown for two weeks in human feeding media (HFM), iCAF$^{ENR}$ or myCAF$^{ENR}$-CM. Scale bar = 50 µm. *p≤0.05 **p≤0.01

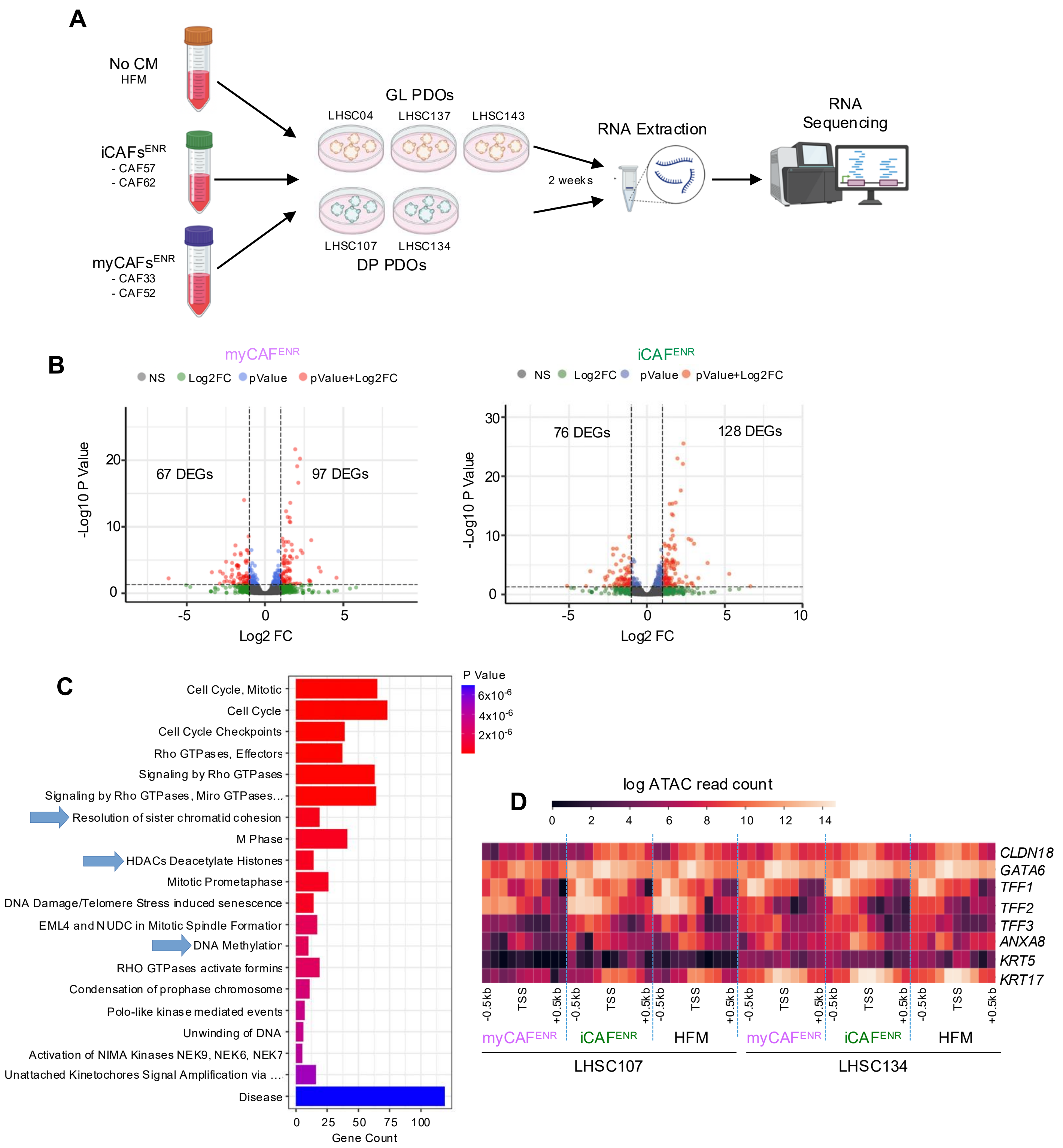


**Supplementary Figure 7 (Jaune-Pons et al.,2026). Molecular analysis of PDOs treated with CAF-conditioned media**. **(A)** Experimental scheme. Five PDOs were grown for two weeks in human feeding media (HFM), $iCAF^{ENR}$-CM, or $myCAF^{ENR}$-CM. After treatment, RNA-seq was performed. For computational analysis, GL-PDOs and DP-PDOs were grouped together. **(B)** Volcano Plot showing differentially expressed genes (DEGs) in DP-PDOs between HFM and $iCAF^{ENR}$ (right) or $myCAF^{ENR}$-CM (left) treatment. **(C)** Reactome pathway analysis on GL-PDOs using DEGs between HFM and $iCAF^{ENR}$-CM treatment. Pathways involved in epigenetic reprogramming indicated with blue arrows. **(D)** Focused ATAC-seq showing read counts from -0.5 to +0.5 kb from the TSS of genes associated with classical (*CLDN18, GATA6, TFF1-3)* or basal-like (*ANXA8*, *KRT5* and *KRT17*) pancreatic cancer in LHSC107 and LHSC134 grown for 2 weeks in HFM, $iCAF^{ENR}$ or $myCAF^{ENR}$-CM.

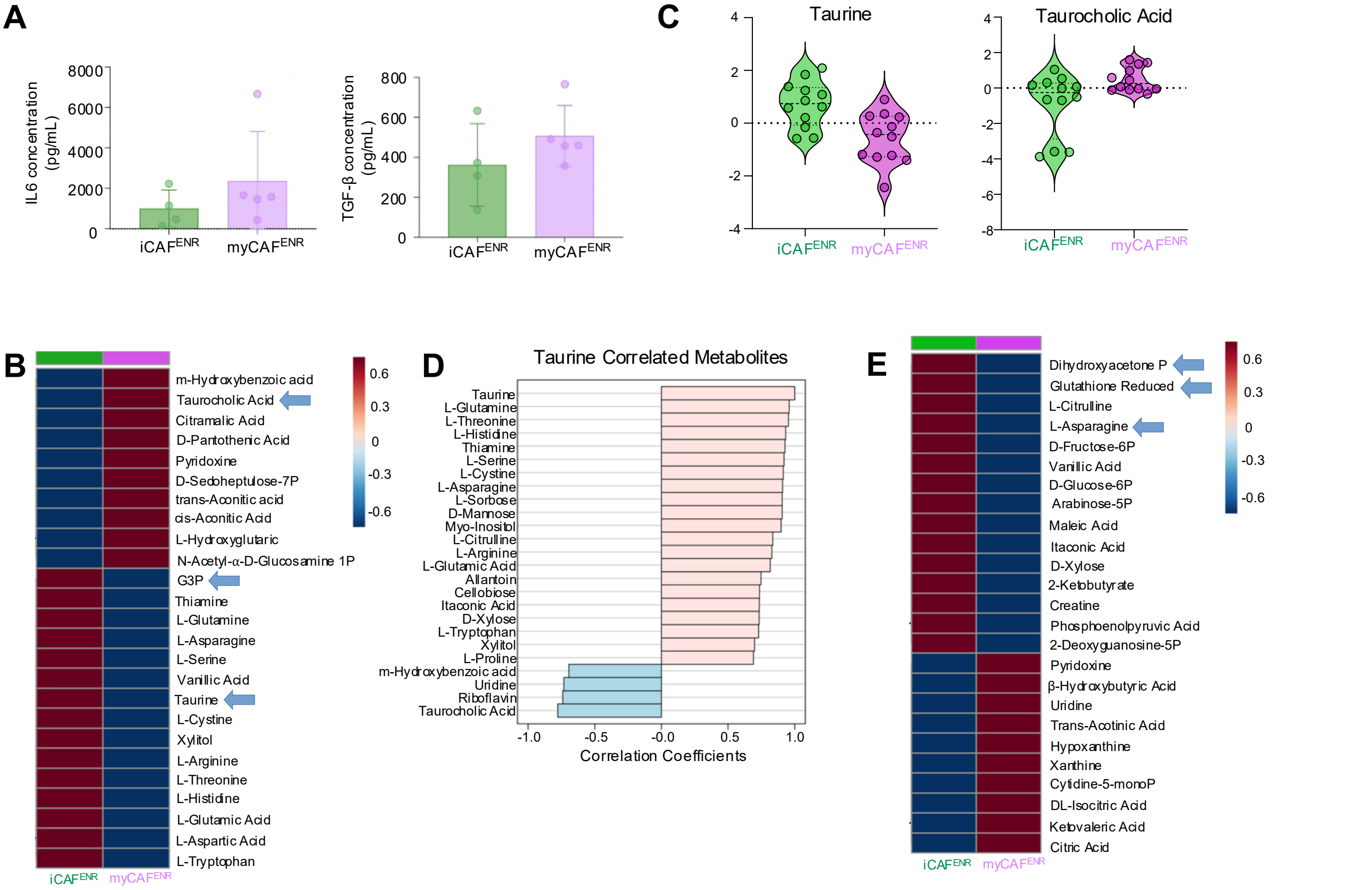


**Supplementary Figure 8 (Jaune-Pons et al.2026). Analysis of CAF-conditioned media. (A)** ELISA assay on CAF supernatants show no significant changes in IL-6 and TGF-β concentrations between iCAF$^{ENR}$ and myCAF$^{ENR}$ conditioned media. **(B)** Mean accumulation of the top 25 metabolites in iCAF$^{ENR}$ and myCAF$^{ENR}$ primary supernatants. G3P and taurine accumulate to higher levels in iCAF$^{ENR}$ vs myCAF$^{ENR}$ samples and taurocholic acid as decreased in iCAF$^{ENR}$ vs myCAF$^{ENR}$. **(C)** Accumulation of taurine (iCAF$^{ENR}$ SEM +/- 0.2524, myCAF$^{ENR}$ SEM +/- 0.2766) and taurocholic acid (iCAF$^{ENR}$ SEM +/- 0.5064, myCAF$^{ENR}$ SEM +/- 0.1990) in iCAF$^{ENR}$ and myCAF$^{ENR}$ supernatants. **(D)** Top 25 metabolites correlated with increased taurine accumulation in iCAF$^{ENR}$ supernatants. Results show the strongest negative correlation is between taurine and taurocholic aaacid. **(E)** Mean expression of the top 25 intracellular metabolites between iCAF$^{ENR}$ (3 samples in triplicate) and myCAF$^{ENR}$ (4 samples in triplicates) samples. This results identify Glutathione Reduced, L-Citrulline and L-Asparagine as overexpressed in iCAF$^{ENR}$ vs myCAF$^{ENR}$ samples.
